# The physics of liquid-to-solid transitions in multi-domain protein condensates

**DOI:** 10.1101/2021.11.15.468745

**Authors:** Srivastav Ranganathan, Eugene Shakhnovich

## Abstract

Many RNA-binding proteins (RBPs) that assemble into membraneless organelles, have a common architecture including disordered prion-like domain (PLD) and folded RNA-binding domain (RBD). An enrichment of PLD within the condensed phase gives rise to formation, on longer time scales, amyloid-like fibrils (aging). In this study, we employ coarse-grained Langevin dynamics simulations to explore the physical basis for the structural diversity in condensed phases of multi-domain RBPs. We discovered a highly cooperative first order transition between disordered structures and an ordered phase whereby chains of PLD organize in fibrils with high nematic orientational order. An interplay between homo-domain (PLD-PLD) and hetero-domain (PLD-RBD) interactions results in variety of structures with distinct spatial architectures. Interestingly, the different structural phases also exhibit vastly different intra-cluster dynamics of proteins, with diffusion coefficients 5 (disordered structures) to 50 times (ordered structures) lower than that of the dilute phase. Cooperativity of this liquid-solid transition makes fibril formation highly malleable to mutations or post-translational modifications. Our results provide a mechanistic understanding of how multi-domain RBPs could form assemblies with distinct structural and material properties.

**Significance Statement:** Assembly of proteins and nucleic acids into dense, liquid-like pockets is associated with several key functions including stress response, gene-regulation, DNA-repair and RNA processing. Several RNA binding proteins such as FUS are known to form liquid-like condensates that progressively harden into more dynamically, solid-like structures, a phenomenon that gets accelerated by disease mutations. In this study, we discover the mechanistic origins of this transition and show that small mutational or posttranslational modifications could result in sharp disorder-order transitions that could characterize accelerated liquid-solid transition in disease mutants.

## 1 Introduction

Spatial segregation of biomolecules is a vital mechanism by which cellular machinery and biochemical reactions get spatio-temporally localized ^1,2^. While classical examples of such spatiotemporal segregation involves physical compartments bound by a membrane, more recent evidence points towards the presence of biomolecule-rich compartments that form and disassemble in response to external cues^3–5^. These structures, known as membraneless-organelles (MLOs) have been associated with diverse cellular functions, ranging from cell division, ribosome synthesis to stress response^3,6–9^. Despite the diversity in function, the physical mechanisms resulting in their formation shows remarkable similarity. The hallmark of these structures is the ability to assemble into droplet-like assemblies via a mechanism called as liquid-liquid phase separation (LLPS)^9–11^.

A key class of membraneless assemblies are ribonucleoprotein granules (RNPs) which are a heterogenous mix of RNA-binding proteins (RBPs) and RNA ^12–14^. Proteins which are localized within RNP granules are often RBPs rich in low-complexity (LC) domains enriched in a select few amino acids ^3,14–16^. The LC-domains, or prion-like domains (PLD) typically display a high degree of intrinsic disorder ^3,14–16^. These IDRs have been previous reported to be key drivers of phase separation in RBP into liquid-like droplets. The interaction of RBPs with nucleic-acids, is mediated by the folded domains localized within the RNA-binding domain (RBD) of the proteins – the RNA-recognition motif (RRM) and the zinc-finger domain (ZnF)^14^. The multi-domain architecture of these proteins also results in multiple modes of inter-molecular interactions (PLD-PLD and PLD-RBD) stabilizing the droplet phase^17^. In their seminal work involving the FUS family of proteins, Wang et al demonstrated that phase separation could be driven by hydrophobic interactions involving the PLD or through pi-pi and pi-cation interactions between the PLD and the RBD^14^. A switch in the predominant mode of inter-molecular interaction (PLD-PLD vs PLD-RBD) not only results in altered threshold concentrations for phase-separation but also structures with different material or dynamic properties^14^. The PLD of FUS is known to form ordered amyloid-fibrillar structures^18^. On the other hand, while full length FUS proteins phase separate into spherical droplets with liquid-like properties initially^19^, they are known to progressively reorganize and assume more filamentous geometries which are dynamically solid-like^20^. This liquid-solid transition gains further importance in disease biology where mutations to the FUS LC domain are known to accelerate the process^21^. Several experimental studies have probed the ability of FUS, FUS-PLD and FUS variants to spontaneously self-assemble into liquid-like droplets in vitro^14,18,22–24^. Structural elucidation of FUS assemblies using solution and solid-state NMR studies have revealed that the FUS PLD can assume a disordered configuration (in liquid-like FUS droplets)^19^ or an ordered solid-like amyloid cross-*β* fold in FUS hydrogels^18^. The wide range of structural, and dynamic phases accessed by FUS makes it a useful model protein to study structural (and dynamic) transitions in multi-domain RNA-binding protein assemblies.

Biomolecular simulations of varying resolutions have previously been employed to study the equilibrium^22,25^ and dynamic properties (droplet surface tension, shear viscosities)^22^ of FUS droplets. Simulations have also been employed to identify the key inter-molecular interactions stabilizing the condensed phase in FUS droplets^26^. A recent computational study by Garaizar et al. demonstrates how an increase in strength of PLD-PLD interactions could result in spatially heterogenous condensates^27^. While the thermodynamics of FUS phase separation has been extensively characterized using computational models, the factors influencing the morphology of the assemblies is yet to be systematically explored. Further, while most of the earlier computational models can capture the spherical droplet phase in FUS, the ‘fibril-like’ ordered state hasn’t been captured in simulations.

In this study, we employ a coarse-grained (CG) representation of the full-length FUS protein to understand the physical mechanisms underlying emergence of order in multi-domain protein assemblies such as those constituted by RBPs. Using this CG representation, we explore how the variation of homo-domain (PLD-PLD) and hetero-domain (PLD-RBD) interaction strengths within a narrow, biologically relevant regime (< 1 kT) could give rise to self-assembled structures with diverse morphologies. We scan a broad range of conditions to establish a relationship between interaction networks, phosphorylation, intra-cluster density and structural order within the condensed phase. Our simulations show that biomolecular assemblies could transition from a disordered to an ordered phase by switching the predominant mode of interaction stabilizing the condensed phase. More generally we observe the emergence of an ordered phase in a first-order like phase transition where loss of entropy due to local ordering of individual polymer chains in fibrils is compensated by favorable interaction energy in a more dense ordered phase. Our simulations also show that the disorder-order structural transitions are also accompanied by dramatically altered intra-cluster dynamics of polymers making them biologically relevant.

## Model

FUS proteins contain two domains critical for their phase separation into condensates – a low complexity, prion-like domain (PLD), and an RNA-binding domain. The prion-like domain and the RGG region (arginine, glycine rich region) of the RBD exhibit a high degree of disorder^28^. Earlier studies have established the ability of the full length FUS proteins^14^ as well as the prion-like domains to form liquid-like droplets^14^, at different saturation concentrations.

### Coarse-graining resolution

In this work, we employ a coarse-grained model for a model scaffold protein, FUS where we coarse-grain stretches of 8 amino acids on the PLD into an effective interaction site (Fig.1). In effect, the 160-residue long PLD of FUS is modeled through 20 effective interacting beads. The RGG (arginine-glycine rich) region is represented at a resolution of 5 amino-acids per bead, with 20 effective RGG interaction beads making up the coarse-grained FUS chain. The choice of different ratios of residues per bead was to ensure that all regions in the coarse-grained polymer chains have a comparable contour length of 20 beads each, thereby ensuring that the orientational ordering is not due to a difference in contour lengths. The slight difference in ratio of amino acids/bead allows us to study ordering due to enthalpic effects and rule out effects due to differences in local rigidity. The charges of the PLD and the RBD beads were fixed based on the net charge of an amino acid stretch coarse-grained by the bead (see Supplementary Table S1 for the FUS sequence). The net charge on the RBD is positive due to high density of arginine residues in the RGG. In order to model phosphorylated PLD (Fig.1, phosphorylated FUS), we assume 31 potential phosphosites^29,30^ (Supplementary Table S2). For simulations involving RNA chains, we employed semi-flexible polymer chains mimicking 200-residue long, unstructured single-stranded, homopolymer RNA. Each bead of the coarse-grained RNA chain represents a single RNA base and is negatively charged.

**Figure 1:**
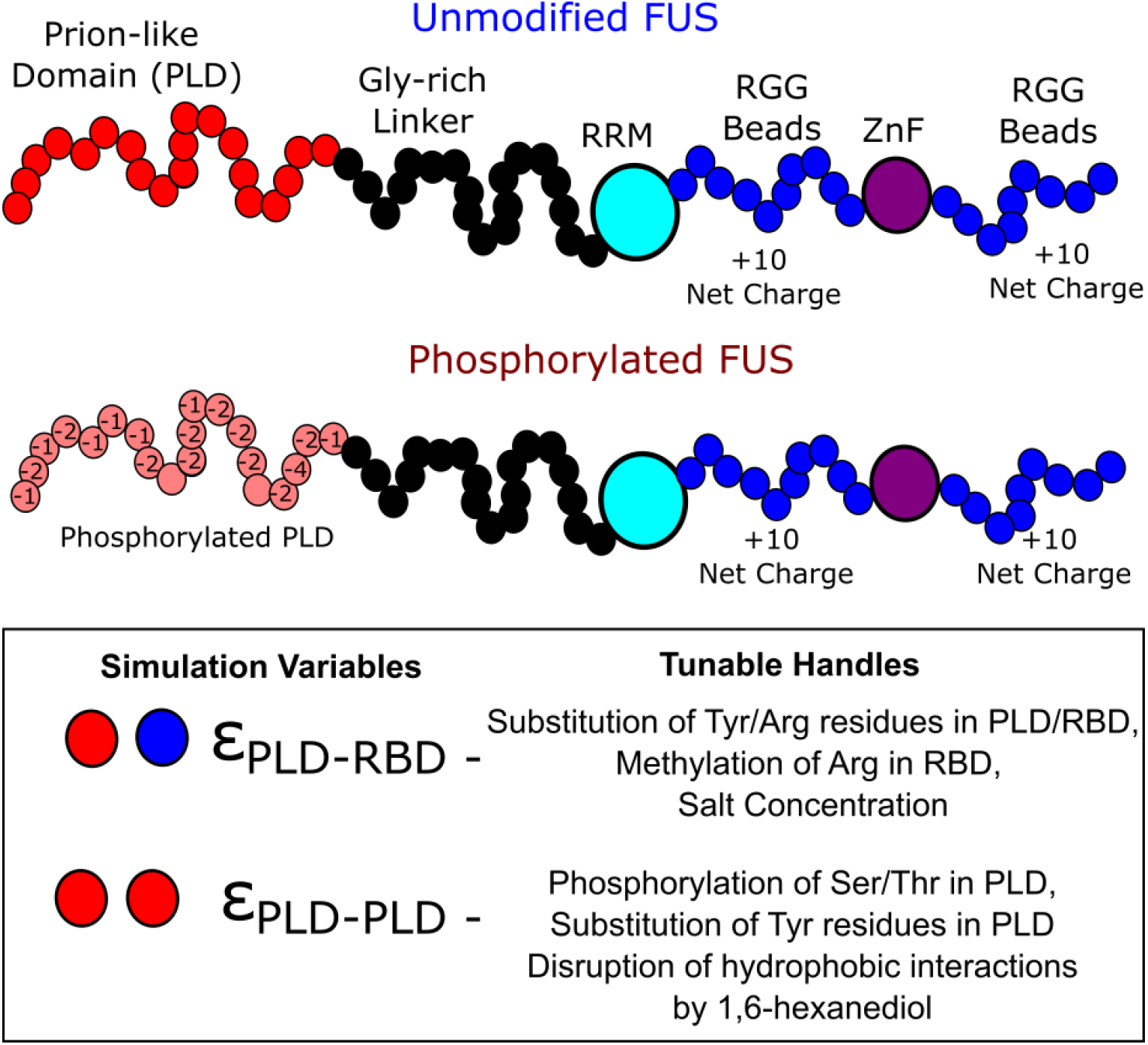
The Model. Coarse-grained representation of unmodified and fully phosphorylated variants of FUS. Red beads represent 8-amino acid long coarse-grained PLD stretches. Blue beads represent 5 amino-acid long stretches from the RGG region of FUS. The glycine rich linkers are shown in black. Two folded domains, the RNA-recognition motif (RRM) and the Zinc Finger Domain are modeled as large idealised beads. The two key simulation variables and their physical in this study, *ε*_*PLD*−*PLD*_ and *ε*_*PLD*−*RBD*_ are shown in the box. In Supplementary Table.S1 and S2, we provide the FUS protein sequence, and the location of phosphorylation sites used for assigning charges to the coarse-grained beads of the PLD and RGG. Note that this schematic is for representative purposes and not drawn to scale.

### Size of coarse-grained beads

The folded stretches of the protein, RRM and the ZnF motif, are modeled as large effective interaction domains. The size (radius) of the RGG and PLD beads was set to 6 Angstroms while the folded RRM and ZnF motifs were of 8 Angstroms. For bead size calculations, we assume that the 6-8 amino-acid stretches within the PLD and RGG behave like random coils. According to the Flory theory, polymer chains scales by N_aa_^γ^σ_aa._ For random coil regions of the protein (PLD region, Linker region and RGG region of RBD), the scaling exponent of ?=2/5 was used while the compactness of the folded ZnF and RRM domains was modeled with a ? of 0.3. We assume a σ_aa_ of 4.5 A for each amino acid. Based on these values, we get a bead radius of about 6 Angstrom (per 6-8 residues) for disordered regions of the protein and ∼8 A for the folded domains. In Supplementary Table S3, we provide details of the different model parameters chosen for the study. The RBD and the PLD are connected by a flexible linker which does not participate in interactions, mimicking the glycine-rich region in FUS. The glycine-rich linker (∼100 residues long) is modeled with 20 beads in our study. Each effective interaction site harbors a charge based on the corresponding region of the full-length protein sequence, and based on the phosphorylation state of the protein. The persistence length (*L*_*p*_) of the PLD and RGG regions was set to 4 residues while that of the glycine linker was set to 1 residue to model greater flexibility of the Gly-rich region.

### Rationale behind choice of key simulation variables

The RRM and the ZnF regions of the protein are folded domains which are primarily involved in FUS-RNA interactions^31,32^. These regions have not been reported to play a role in the self-assembly of FUS. The low-complexity PLD and the RGG regions of the protein, on the other hand, have been established to be the primary drivers of LLPS in experimental studies involving FUS^17,32^. Therefore, in this study, we primarily focus on the interplay between PLD-RBD and PLD-PLD interactions, assuming that the RRM and ZnF domains do not play a key role in self-assembly. Therefore, we use two interaction variables -- *ε*_*PLD*−*PLD*_ and *ε*_*PLD*−*RBD*_ – as effective interaction parameters (see Methods) that capture the physicochemical properties of the two FUS domains (PLD and RBD) as well as solvent conditions. Physically, any change in solvent conditions or in the protein sequence would manifest itself via a change in either (or both) these interaction parameters (Tunable handles in Fig.1). Therefore, for the rest of this paper, we present phase diagrams wherein we use *ε*_*PLD*−*PLD*_ and *ε*_*PLD*−*RBD*_ as our key sequence and environment-dependent phase parameters.

## Methods

Coarse-grained models for proteins/peptides, at varying resolutions, have been used to study relationship between biomolecular structure and self-assembly^25,33,34^. Due to the fewer degrees of freedom and higher computational tractability, these coarse-grained models are efficient tools to probe structural transitions in self-assembling polymers^35^. In order to study the diversity in self-assembled states in response to varying PLD-PLD and PLD-RBD interaction strengths, we performed Langevin dynamics simulations. To simulate self-assembly, we perform simulations with 100 coarse-grained FUS chains (61-beads each) in a cubic box with periodic boundary conditions. Previous computational studies with FUS have established that dynamic and equilibrium quantities converge at system sizes of 50 and above, suggesting that a system size of 100 monomers is large enough to avoid finite size effects^22,25^. The coarse-grained FUS polymer beads interact via the following potential functions. The neighboring beads in a polymer chain are bonded via the harmonic potential with energy

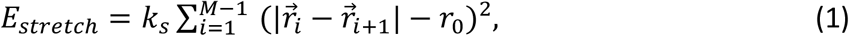

where 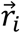 and 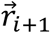 correspond to the positions of the *i*^*th*^ and (*i* + 1)^*th*^ beads, respectively; *r*_0_ is the resting bond length. *k*_*s*_ denotes the spring constant and has units of kT/Å^2^ and was set to 5 kT/ Å^2^. This interaction ensures the connectivity between the adjacent beads of the same polymer chain.

To model semi-flexible polymers, neighboring bonds in a polymer chain interact via a bending potential

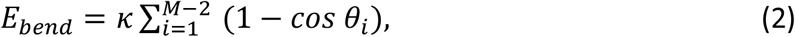

where *θ*_*i*_ refers to the angle between *i*^*th*^ and (*i* + 1)^*th*^ bond. Here, *κ* the bending stiffness, has units energy, kT. The value of *κ* was chosen such that the persistence length (Lp = *κ*/*kT*) of the polymer chain was ∼4 FUS PLD/RGG/Linker beads. It must therefore be noted that the contour length of the PLD (L_PLD_) and the RBD (L_RBD_) regions in the coarse grained polymer are 5 times that of the persistence length of the polymer. The contour length of the full length coarse-grained FUS is 15 times that of the persistence length, indicating the semiflexible nature of the modeled polymer chains. The persistence length of RNA chains was set at 8 beads, with a contour length of 200 for the RNA chains.

Interactions between any pair of non-bonded beads were modeled using the Lennard-Jones potential.

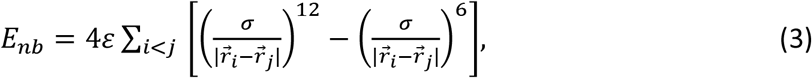

for all 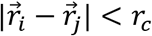. Here *r*_*c*_ refers to the interaction cutoff beyond which LJ interactions are neglected. The cutoff for LJ calculations were set to 2.5 times of *σ*. The values of *ε* and *σ* corresponding to different pair-wise interactions and bead types can be found in Supplementary Table S3.

To model electrostatic interactions, we use the standard Debye-Huckel electrostatic potential which has the form,

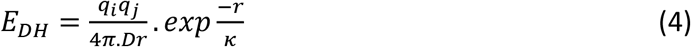

*q*_*i*_ and *q*_*j*_ are the charges of the two interacting beads. Each RNA bead carries a single negative charge in our simulations. As previously demonstrated by Dignon et al.^25,36^, the Debye-Huckel potential can robustly capture features of phase separating proteins such as TDP43 and FUS. We therefore use the Debye-Huckel potential with a screening length of 1 nm (corresponds to an ionic strength of 1 mM) in our simulations to mimic the screening of electrostatic interactions in a cellular environment.

The attractive part of the LJ potential, *ε*, is tuned to vary the strength of the interaction. In this study, the strength of interaction between PLD beads is referred to as *ε*_*PLD*−*PLD*_ while PLD and the disordered regions of RBD (RGG beads) interact with a strength *ε*_*PLD*−*RBD*_. The pairwise interaction strengths for all bead types – PLD, RBD (refering to the disordered parts of RBD), linker, RRM and ZnF domains are listed in Supplementary Table S3. We used the LAMMPS molecular dynamics package^37^ to perform the simulations, where the simulator solves Newton’s equations with viscous force, and a Langevin thermostat ensuring an NVT ensemble with temperature *T =* 310 K. An integration timestep (*dt*) of 20 fs was used for the simulations.

### Metadynamics

To enable efficient sampling of the self-assembly landscape and to ensure the formation of a single large cluster, we employed metadynamics^38^. In this method, the evolution of the self-assembling system is influenced by a history-dependent biasing potential. Energy gaussians are deposited during the course of a trajectory along the free energy space for a chosen reaction coordinate that effectively describes the process of self-assembly. The deposited potentials are then used to reconstruct the free energy profiles along this chosen reaction coordinate^38^. In order to drive self-assembly and observe coalescence of protein-rich clusters into a single large cluster, we used the radius of gyration of the peptide system, 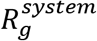 as the reaction coordinate. 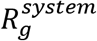 is the radius of gyration of an imaginary polymer that links together the center of masses of the individual polymer chains in the simulation. Large values of 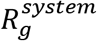 correspond to a fully mixed system whereas smaller values indicate a self-assembled state (Supplementary Fig.S1 and S2). The systems were simulated till the free energy profiles showed convergence.

### Simulation Protocol

We first run metadynamics simulations (with the 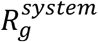 as the collective variable) to compute the free energy profiles and also ensure that the coarse-grained chains coalesce into one large cluster at simulation timescales. This single large, coalesced structure corresponding to the free energy minima (R_g_^system^ at ∼50 A in Supplementary Fig.S1) was then used as a starting configuration for conventional Langevin dynamics trajectories (with no biasing potential) of at least 10 microseconds or till the simulations reached convergence. In Supplementary Fig.S3 and S4, we provide proof of principle for convergence of our simulations. To ensure convergence, we also start our simulations with different initial configurations -- fully mixed disordered state, fibril-like core state -- and then allow these structures to evolve under different regimes of ε_PLD-PLD_ and ε_PLD-RBD_. The computed order parameters were found to be independent of the starting configuration, thereby establishing convergence (Supplementary Fig.S4). These unbiased 10-microsecond long simulation trajectories were used to compute statistical quantities (orientational order, inter-protein contacts. These unbiased trajectories were also used to compute intra-cluster dynamics and autocorrelation functions. We run 10 independent Langevin dynamics trajectories (Supplementary S3).

### Orientational Order Parameter

To characterize structural order in the FUS assemblies, we employ the nematic order parameter which is commonly used in liquid-crystal literature to quantify the extent of alignment of molecules along a reference axis^39–41^. We first define the long axis of each of the ‘*N*’ FUS chains within the self-assembly by a unit vector 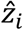 which connects the two terminal beads of the PLD domain. We compute the extent of alignment in each local region around the ‘N’ polymer chains using the local nematic order parameter S.

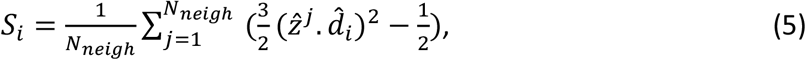

where the index ‘i’ ranges from 1 to N, the total number of polymer chains in the system.

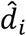 in the above equation is a unit vector that corresponds to the direction of local alignment for a set of N_neigh_ polymer chains (including the i^th^ chain) that define the local region around any polymer chain ‘i’. To define a local region around each polymer chain, any chain that lies within an interation radius of 2.5*σ* is considered to be in physical contact. Therefore, using equation 5 we get ‘N’ such scalar order parameter values, each corresponding to a local region defined by every polymer chain in the system. Next, we average over all local order parameters:

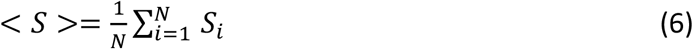

In our study, we use <S> (computed using Eqn.6) to define the extent of local ordering within a self-assembled structure. In the rest of this particle, we use S instead of <S> to refer to the mean orientational order in the system. This order parameter S ranges from 0 to 1. *S* → 0 indicates weak orientational order, while *S* → 1 suggests high local orientational order.

In order to obtain the director vector, 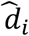required to compute the orientational order parameter using Eqn.5, we define a second-order tensor, 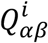 which reflects nematic ordering for each local region corresponding to the ‘*N*’ FUS chains in the system.

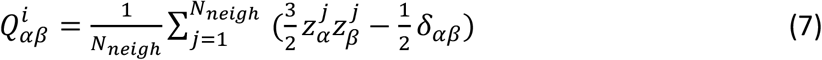

where 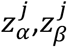 are component vectors (*α, β =*x,y or z) corresponding to each of the *N*_*neigh*_ chains (including the i^th^ chain) that the *i*^*th*^ chain is physically in contact with. The index *i* refers to the identity of each chain in the system. We therefore have *N* such independent local order tensors 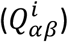 -- each one corresponding to the region defined by every FUS chain and its corresponding N_neigh_ neighbors. From Eqn.7, we obtain 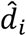 by computing the eigenvector corresponding to the largest eigenvalue of the tensor 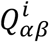. We therefore have N such eigenvectors, 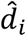, each defining the direction of alignment in the region around the i^th^ chain. These director vectors are used to compute local orientational order S_i_ using equation 5.

## Results

### Disorder-order transition is an outcome of hetero to homo-domain interaction network switching

FUS is known to spontaneously phase separate in vitro and also act as a scaffold protein in RNA-protein condensates in the cell^15,19^. Previous studies have shown that FUS droplets are stabilized by two primary modes of interactions – PLD-PLD interactions and PLD-RBD interactions. Here, we systematically vary the strength of these two interactions -- *ε*_*PLD*−*PLD*_ & *ε*_*PLD*−*RBD*_ – to study how it influences the nature of the FUS condensate.

In order to drive the self-assembly of the proteins into a single, large, condensed phase at simulation timescales, we employ metadynamics simulations. We use the radius of gyration of the system 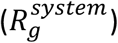 of 100 protein chains as a collective variable to define the state of the system. The free energy profiles were computed along this reaction coordinate, with minima around small values of 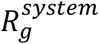 indicating a fully assembled equilibrium state. In Supplementary Fig.S1, we demonstrate the free energy profiles for four such scenarios, with varying strengths of hetero-domain (*ε*_*PLD*−*RBD*_) and homodomain (*ε*_*PLD*−*PLD*_) interactions. The free energy profiles suggest that, for the protein concentration chosen for the study (200 uM) the self-assembled state is favored even for relatively weak interaction strengths (*ε*_*PLD*−*RBD*_ of 0.15 kT && *ε*_*PLD*−*PLD*_ of 0.3 kT).

Upon establishing that the condensed phase indeed corresponds to an equlibrium state, we then proceed to characterize the interaction network of the condensed phase, for a broad range of interaction parameters. In Supplementary Fig.S5, we plot the number of inter-molecular contacts per chain as a function of strengths of homotypic interactions between amino acids in the PLD domain and heterotypic interactions between PLD-RBD domains, *ε*_*PLD*−*PLD*_ and *ε*_*PLD*−*RBD*_, respectively. For weak interaction strengths, the interaction network within the FUS cluster is sparse (*Qinter* <*=* 100 inter-molecular contacts per chain in Supplementary Fig.S5). For strong PLD-PLD interactions, we observe a dense inter-molecular contact network (> 100 contacts per chain in Supplementary Fig.S5) suggesting a different intra-cluster interaction network in this regime.

Upon establishing variation in the extent of intermolecular connectivity within the droplet upon tuning the interaction parameters, we probe the nature of interactions that make up these intermolecular contacts. In Fig.2A, we plot the ratio of PLD-PLD to PLD-RBD intermolecular contacts to elucidate the dominant intermolecular interactions that stabilize the condensed phase. For weak *ε*_*PLD*−*RBD*_ and strong *ε*_*PLD*−*PLD*,_ the FUS clusters are stabilized by homotypic interactions between PLD domains from different chains. On the other hand, in the limit of weak *ε*_*PLD*−*PLD*_ and strong *ε*_*PLD*−*RBD*_, hetero-domain interactions dominate. In a narrow range of interaction strengths, the contact ratios → 1 (*Q*_*PLD*−*RBD*_ *= Q*_*PLD*−*PLD*_). In this limit, the intra-cluster environment exhibits both homo- and hetero-domain contacts with equal likelihood.

Tuning the strength of homo- and hetero-domain interactions in the thermally relevant regime results in structures that are stabilized by distinct modes of interaction (Fig 2A and Supplementary Fig.S5). Wang et al have previously established that a hierarchy of interactions – homo- and hetero-domain – can drive phase-separation of FUS into protein-rich droplets^14^. We, therefore, probe whether a switch in the primary mode of inter-molecular interactions upon varying the interaction strengths could result in altered structural properties of the assembly. To characterize the morphology of the assemblies, we employ the nematic orientational order parameter *S* which defines the extent of local orientational order for the prion like domains within a cluster. This order parameter, *S* can take any value from 0 to 1. *S* → 1 indicates a high degree of orientational order. On the other hand, *S* → 0 suggests an entirely disordered system. Using this order parameter, we capture the extent of orientational order as a function of varying interaction parameters *ε*_*PLD*−*PLD*_ and *ε*_*PLD*−*RBD*_. In Fig.2B, we plot *S* as a function of increasing PLD-PLD interactions for two different strengths of PLD-RBD interactions. Interestingly, an increase in *ε*_*PLD*−*PLD*_ results in a first order-like, disorder → order transition for weak *ε*_*PLD*−*RBD*_. A further affirmation of the discontinuous nature of this transition could be ascertained from the evolution of the distribution of S for increasing values of (Fig.2C). For weak homodomain interactions (*ε*_*PLD*−*PLD*_<= 0.5 kT in Fig.3B), we see the distributions peaking at low values of S (∼0.2), suggesting that the assemblies never access the ordered states in this interaction regime. Interestingly, an increase in *ε*_*PLD*−*PLD*_ from 0.2 to 0.5 kT results in a negligible shift in distribution. However, a slight increase in homodomain interaction strength from 0.5 to 0.512 kT results in a shift from a unimodal distribution (fully disordered) to a bimodal distribution with peaks corresponding to both the ordered and disordered states. Such a bimodality is a typical signature of discontinuity. A further increase in interaction strength to 0.6 kT results in a unimodal distribution with the assemblies populating the ordered state alone. For stronger *ε*_*PLD*−*RBD*_ interaction, the S vs *ε*_*PLD*−*PLD*_ curve saturates at lower values of the order parameter, owing to a competition between homo-domain and heterodomain interactions.

**Figure 2:**
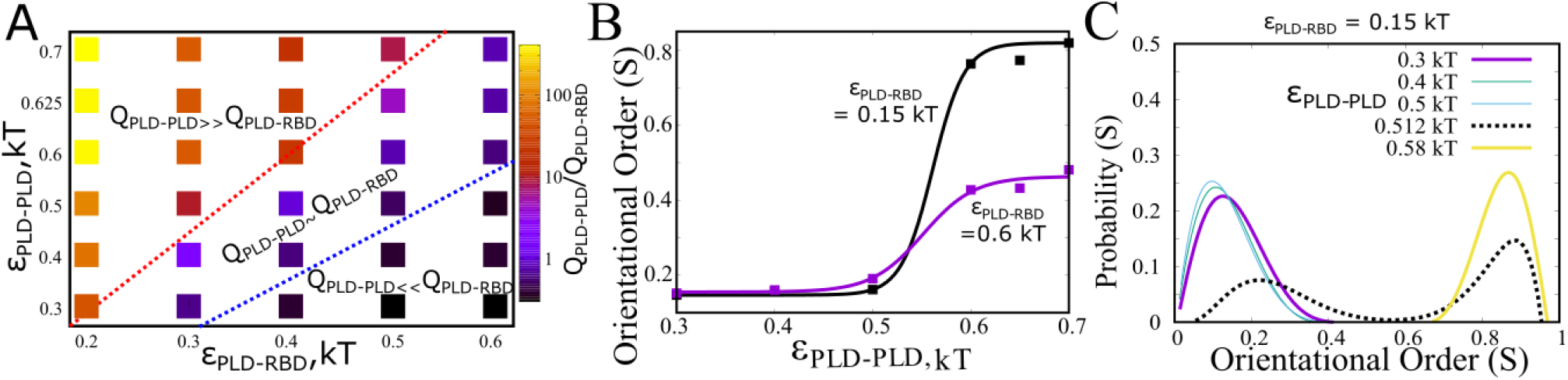
Structural Transitions at different regimes of *ε*_*PLD*−*PLD*_ & *ε*_*PLD*−*RBD*_. A) Switching of predominant interaction network stabilizing the condensate, as a function of varying *ε*_*PLD*−*PLD*_ & *ε*_*PLD*−*RBD*_. *Q*_*PLD*−*PLD*_ and *Q*_*PLD*−*RBD*_ refer to the inter-molecular contacts per chain between PLD-PLD and PLD-RBD domains, respectively. B) Local orientational order as a function of *ε*_*PLD*−*PLD*_, for different strengths of *ε*_*PLD*−*RBD*_. The solid curves are just shown as a guide to eye. C) Distribution of the local orientational order parameter (S), for different values of *ε*_*PLD*−*PLD*_. The distributions were computed across 100 different orientational local order parameter values and 50 different structures at equilibrium.

In Fig.3, we show how a variation in these two parameters gives rise to different structural phases with varying degrees of nematic ordering. In the *ε*_*PLD*−*PLD*_-*ε*_*PLD*−*RBD*_ phase diagram in Fig.3, we define all structures with *S* < 0.4 as disordered assemblies. *S* > 0.6 were defined as ordered assemblies while those with intermediate values of *S* were labeled as partially ordered structures. The S values were averaged across 100 different equilibrium configurations, and across 100 polymer chains. When both homo- and heterodomain interactions are weak (*ε*_*PLD*−*PLD*_ & *ε*_*PLD*−*RBD*_ < 0.5 kT), we encounter structures which are disordered, and exhibit spherical geometry. As we increase the strength of homodomain interactions, in the limit of weak heterodomain interactions (*ε*_*PLD*−*RBD*_ < 0.5 kT), we encounter ordered structures with the PLD domains displaying a high degree of orientational order with ‘fibril-like’ geometry. When both interactions are competitively strong (*ε*_*PLD*−*PLD*_ and *ε*_*PLD*−*RBD*_ > 0.5 kT), we observed structures with partial order. Therefore, in the regime where homotypic interactions between PLD domains dominate (*Q*_*PLD*−*PLD*_ ≫ *Q*_*PLD*−*RBD*_ in Fig.2A) the self-assembled state has a high propensity to organize into *ordered* structures stabilized by PLD-PLD contacts. An interplay between these homodomain and heterodomain interactions can therefore result in structures with varying morphologies.

**Figure 3.**
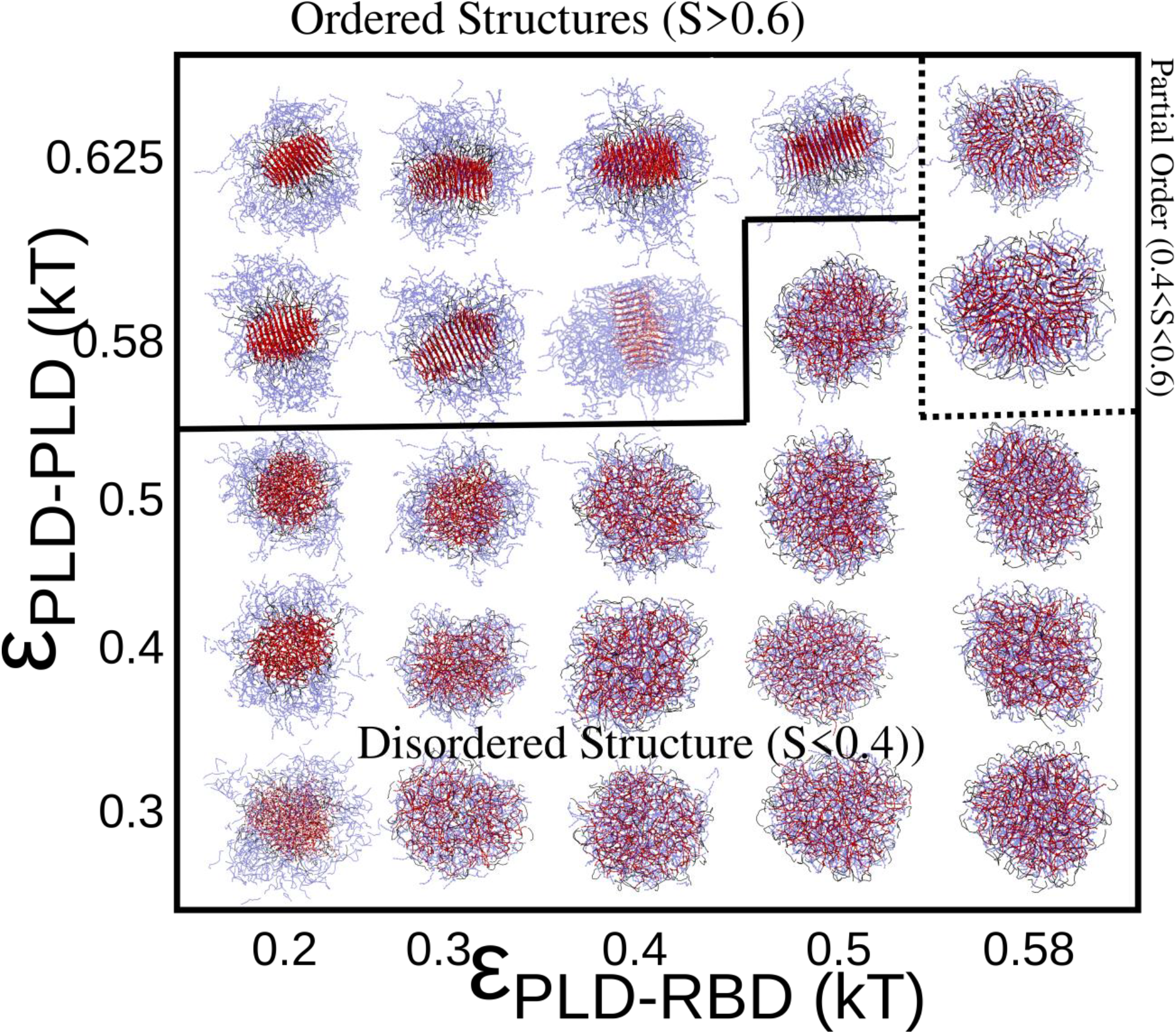
Structural Phases. Phase diagram desribing the various structural phases in response to varying strengths of homo-domain(*ε*_*PLD*−*PLD*_) and heterodomain (*ε*_*PLD*−*RBD*_) interactions. Structures are described as ordered, disordered and partially ordered based on the mean value of the nematic orientational order parameter S. Assemblies with S > 0.6 are labeled as ordered while those with S < 0.4 are labled as disordered structures. PLD residues are shown in red, while RBD regions are shown in blue.

### Phosphorylation of the PLD is a barrier to the formation of ordered structures

Our results so far show that even within a narrow range of interaction strengths, in the thermally-relevant limit, the interplay between homo- and hetero-domain interactions can result in ordered structures. Phase-separated structures in the cell could localize proteins at concentrations as high as 50-100 times that of their average bulk concentrations^42,43^. Such a large local density of prion-like domains in these condensates makes these structures prone to forming filamentous structures. Previous experimental studies suggest that post-translational modifications could potentially act as a barrier to liquid-solid transitions in cellular condensates^29,44^.

In this context, we modeled a key post-translational modification that is typically associated with the FUS PLD, i.e phosphorylation. The FUS PLD in primates harbors 31 potential phoshposites^30,45^. Here, we model phosphorylation by introducing a single negative charge per phosphosite in our coarse-grained PLD beads. The net negative charge on each phosphorylation PLD bead corresponds to the number of phosphosites^30^ residing in a particular 8-residue sequence that the coarse-grained bead maps onto (see Supplementary Table S2). The net charge on the phosphorylated variant of the PLD is -31. In Fig.4A, we compare the propensity of three variants of FUS –the PLD domain only, Full Length FUS, and phosphorylated FUS – to form ordered structures. The PLD control simulations reveal that the PLD domain can undergo a disorder-order transition on its own in the limit of *ε*_*PLD*−*PLD*_ > 0.5 kT. Similarly, the Full Length FUS, for *ε*_*PLD*−*RBD*_ *=* 0.5 kT, also shows a *ε*_*PLD*−*PLD*_-dependent disorder-order transition, with lower < *S* > values than the PLD-only control simulations. The phosphorylated FUS, on the other hand, shows no such propensity to form highly ordered structures (S > 0.6) even at *ε*_*PLD*−*PLD*_ approaching 1 kT. This result suggests that the high local density of negative charges in the PLD of phosphorylated FUS results in an abrogation of order formation in these structures.

**Figure 4:**
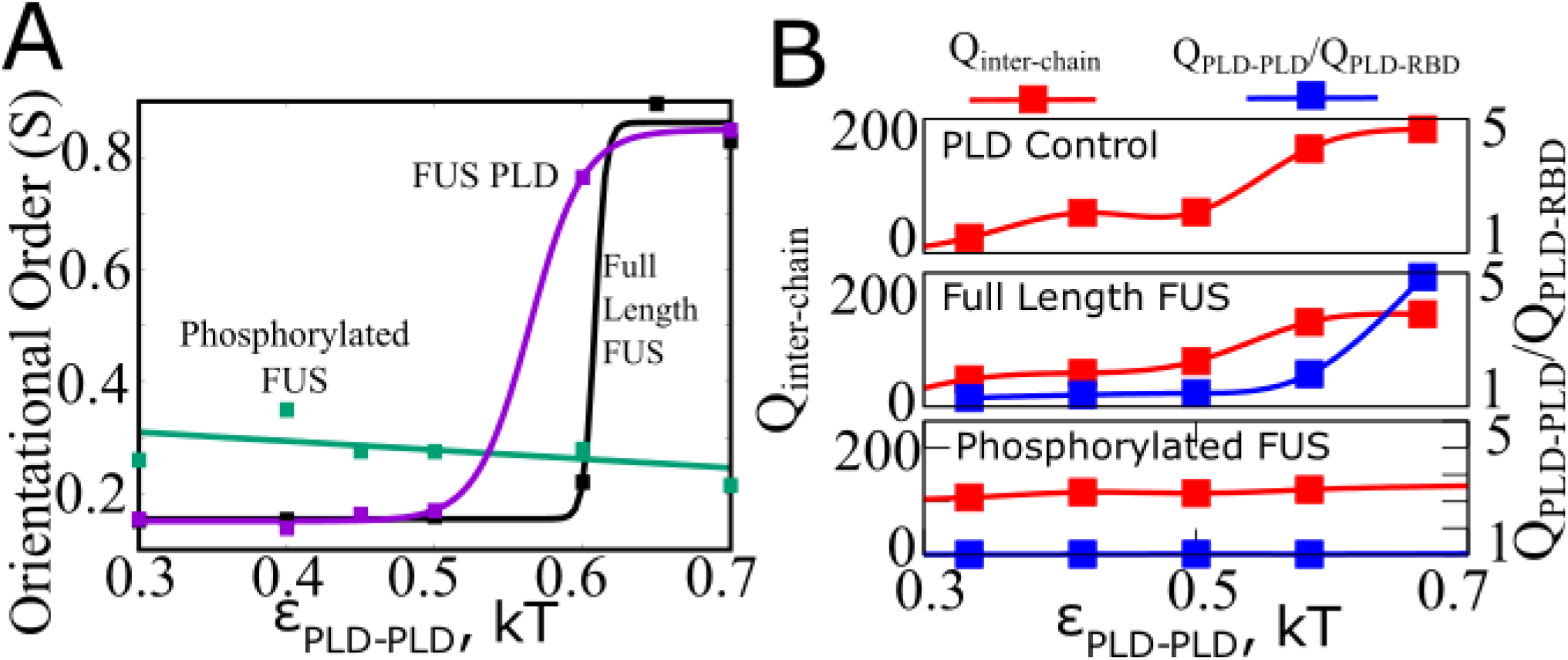
Phosphorylation abrogates order. A) Orientational order as a function of *ε*_*PLD*−*PLD*_ for the PLD alone, full-length FUS and phosphorylated FUS. Phosphorylation results in disordered structures even at strong *ε*_*PLD*−*PLD*_. B) Mean inter-chain contacts per FUS chain (*Q*_*inter*−*chain*_) and the contact ratio (*Q*_*PLD*−*PLD*_/*Q*_*PLD*−*RBD*_) for PLD alone (top), full-length FUS (middle) and phosphorylated FUS (bottom) as a function of *ε*_*PLD*−*PLD*_. Phosphorylation results in lower values of contact ratios. The solid lines are guide to the eye.

We further probe whether this loss of ordering in phosphorylated version of FUS is a result of overall loss of inter-molecular contacts within the FUS-cluster. As evident from the *Q*_*inter*−*chain*_ vs *ε*_*PLD*−*PLD*_ curve for the phosphorylated FUS (Fig.4B, bottom panel), the modification to the PLD results in number of intermolecular contacts (*Q*_*inter*−*chain*_) comparable to the unmodified variant. However, unlike the other two scenarios, the phosphorylated FUS shows no increase in contact ratio (*Q*_*PLD*−*PLD*_/*Q*_*PLD*−*RBD*_) even at strong *ε*_*PLD*−*PLD*_ interactions. However, the phosphorylated FUS clusters are stabilized by interactions between the negatively charged phosphorylated PLD and the positively charged RBD. Phosphorylated structures, therefore, show higher ordering than their unmodified counterparts in the limit of weak PLD-PLD interactions (Fig.4A, *ε*_*PLD*−*PLD*_ < 0.5 kT). This ordering, however, is driven by inter-domain electrostatic interactions and order remains unaltered upon an increase in PLD-PLD interaction strength (Fig.4A, green curve). In the corresponding limit of strong *ε*_*PLD*−*PLD*_, the full length FUS displays contact ratio values approaching 5. This shows that phosphorylation results in an upregulation of the heterodomain interactions over homodomain interactions, and thereby results in a more disordered condensed phase.

### Disorder-order transitions are characterized by sharp density transitions

The transition from droplet-like morphology to a more fibrillar morphology in FUS protein assembly is also accompanied by a loss of dynamicity of the condensate^46^. We therefore probe how tuning *ε*_*PLD*−*PLD*_ and *ε*_*PLD*−*RBD*_ alters local densities of PLD and RBDs within the clusters. In Fig.5A, we plot the pair-correlation function, g(r) for different domains in the condensed phase. The g(r) plotted in Fig.5A is the mean number of atom pairs of any type at a given radial distance, normalized by a pair of ideal gas particles for the same bulk density. The g(r) values in Fig.5 refer to the average number of atom pairs of any type (PLD-PLD, PLD-RBD, RBD-RBD) found at any radial distance (between r and r+dr), normalized by the equivalent values for a system of ideal gas particles of the same bulk density. As the peak value of g(r) → 1, the system exhibits gas-like organization. The higher this peak in g(r), (*g*(*r*)^*peak*^ in Fig.5A), the system exhibits more solid-like organization. Also, the higher the g(r) peak, the denser the packing, and potentially less dynamic the condensed phase.

**Figure 5:**
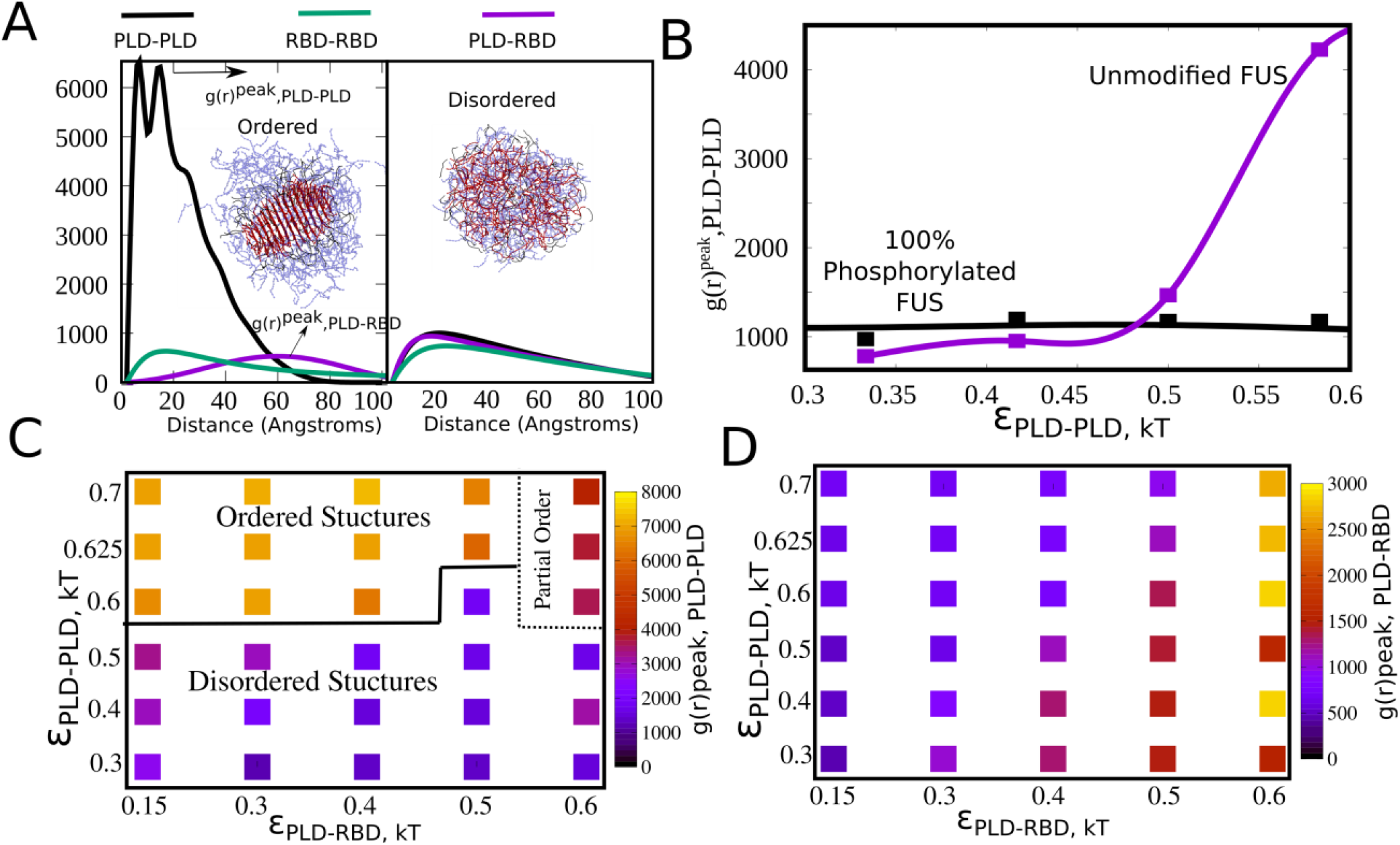
Local Density of Domains. A) Pair correlation function, g(r), for PLD-PLD (black), RBD-RBD (green) and PLD-RBD (purple) contacts within the largest cluster, for ordered and disordered structures. The g(r) here is essentially the probability of finding any two particle types (PLD-PLD, PLD-RBD, RBD-RBD) within a certain distance of each other normalized by corresponding value for a pair of ideal gas particles at the same bulk density. For ordered structures, the g(r) for PLD-PLD interactions peaks several-fold higher than the other two curves suggesting a high local density of PLD domains within the cluster. For disordered structures, the three curves peak at similar values indicating a more homogenous density throughout the cluster. B) The peak values of g(r) plotted as a function of *ε*_*PLD*−*PLD*_. For full-length FUS, the g(r) peak for PLD-PLD contacts shows a sharp sigmoidal transition with increasing *ε*_*PLD*−*PLD*_, suggesting a sharp transition from a low to high density phase. For phosphorylated variants of FUS, such a transition was not observed. C) and D) Phase diagrams showing the density of PLD-PLD and PLD-RBD packing, respectively. An increase in PLD-PLD interactions results in a high packing density for PLD domains within the cluster, as evident from the g(r) peak values for PLD-PLD interactions. In contrast, little variation in PLD-RBD g(r) peaks was obseved throughout the parameter space explored.

In Fig.5A, we show representative g(r) curves for ordered and disordered structures. For ordered structures, the g(r) for PLD-PLD interactions shows peak values several-fold higher than the corresponding curves for PLD-RBD and RBD-RBD interactions. This suggests that the ordered structures have a dense PLD-core and a less dense RBD shell, as also evident from the structures shown in Fig.3. The two peaks in the g(r) profile are indicative of the morphology of the ordered structures. The first peak (∼10 A) in the pair correlation function corresponds to the spacing between adjacent PLD chains (i & i+1th neighbours in Supplementary Fig.S6) in the ordered assembly while the second peak (∼10 A) is indicative of the spacing with the next-nearest (i & i+2th neighbour in Supplementary Fig.S6). The PLDs get packed within a 60 Angstrom radius core within the FUS cluster, and the g(r) curve starts decaying around length scales suggesting that the outer shell of these structures is comprised primarily of the RBD and linker regions. The significantly higher peaks in the g(r) profile for PLD-PLD in Fig.5A is indicative of the difference in packing densities for the demixed, PLD-rich core and the RBD-rich outer shell. For disordered structures formed in the limit of weak interactions (*ε*_*PLD*−*PLD*_ and *ε*_*PLD*−*RBD*_ < 0.5 kT), the g(r) curves for all three pair-wise interactions are comparable, with peak values several-fold lower than the peak g(r) values for PLD-PLD packing in the ordered structures. This result suggests that intra-cluster densities of polymer domains in the disordered structure is more uniform compared to the ordered structures which exhibit a dense core and a less dense shell.

We further probe how phosphorylation influences the peak densities of PLD domains within the FUS clusters by plotting the *g*(*r*)^*peak*^ for PLD-PLD contacts as a function of *ε*_*PLD*−*PLD*_. Strikingly, even at higher *ε*_*PLD*−*PLD*_, the phosphorylated variant shows no change in peak densities within the condensed phase. The full length FUS, on the other hand, shows a discontinuous transition from uniform, less dense structures to the more dense core-shell structures that typify order. We further plot the peak densities for the PLD-PLD and PLD-RBD pairwise contacts as a function of varying *ε*_*PLD*−*PLD*_ and *ε*_*PLD*−*RBD*_. Interestingly, the regime of interaction strengths that corresponds to a high contact ratio (Q_PLD-PLD_ >> Q_PLD-RBD_ in Fig.2A) and high orientational order (Fig.3) also corresponds to large peaks in g(r) for PLD-PLD (Fig.5C). However, the peak values of g(r) for PLD-RBD interactions (Fig.5D) are significantly lower even for strong PLD-RBD interactions.

The g(r) peak values for PLD-RBD show a 5-fold increase as we increase *ε*_*PLD*−*RBD*_ from 0.15 to 0.7 kT. However, the g(r) peak values for PLD-RBD interactions are ∼2-fold lower than those of PLD-PLD interactions, even at strong *ε*_*PLD*−*RBD*_ (Fig.5D). This is consistent with an absence of ordering through strong PLD-RBD interactions. The physical barrier preventing dense packing via PLD-RBD interactions is the high density of charges in the RBD of FUS. While the strength of RBD-RBD interactions remains unaltered across the phase space, an increase in *ε*_*PLD*−*RBD*_ has an indirect effect on the peak densities of RBD-RBD interactions (Supplementary Fig.S7). As we increase strong *ε*_*PLD*−*RBD*_from 0.15 to 0.6 kT, we observe a 3-fold increase in peak heights for RBD-RBD interactions. Therefore, the local densities required for ordered structures cannot be achieved through PLD-RBD interactions, due to the concommitant increase in contact probability of charged RBD-RBD interactions for this packing. Therefore, even for strong PLD-RBD interactions, we observe partially ordered structures instead of the fully ordered structures that are observed at low PLD-RBD and strong PLD-PLD interactions. Therefore, the most feasible mechanism of forming ordered FUS structures is via de-mixing of the PLD and RBD domains within the condensed phase, enabling high packing densities reminiscent of the solid-like state.

### Assemblies can switch from fully-mixed to multi-phase architecture on switching interaction networks

Multi-component mixtures (composed of several protein types) can phase separate into multi-phase assemblies with sub-phases which localize different components based on interaction preferences. Diversity in interactions and interfacial tensions has been known to be a key driver in segregation of components between different sub-phases. Due to the multi-domain architecture of RBPs and the ability to form clusters stabilized by different interaction networks (Fig. 2A), we probed whether multiphase assemblies could result from single component mixtures of FUS alone. In Fig.5 we show how the interplay between the homo-domain and hetero-domain interactions can result in hugely altered local densities of the PLD and RBD domains within the condensates. Given how density differences are a vital signature of multi-phase condensates, we looked at how different domains get packed within the single-component FUS assemblies stabilized by different interaction networks. In Fig.6, we show how FUS condensates could exhibit different sub-cluster organization for different combinations of interaction strengths. For weak interactions (*ε*_*PLD*−*PLD*_ and *ε*_*PLD*−*RBD*_ < 0.5 kT), when PLD-PLD interactions dominate (*ε*_*PLD*−*PLD*_ = 0.4 kT and *ε*_*PLD*−*RBD*_ = 0.2 kT), we observe spherical clusters with a multi-phase structure. Here, PLD segments form the core of the condensates while the RBD domains organize themselves in the outer shell. This is evident from the distributions of the radius of gyration of PLD domains which peak at a much lower value (∼35 A), compared to that of RBD domains (∼70 A) showing that the PLD domains form a condensed core. However, these core-shell structures formed by PLD, in the limit of weak interactions, do not exhibit nematic order. As the strengths of the two interactions become comparable, we observe single-phase clusters where the two domains are fully mixed (Fig. 6B), with the radius of gyration distributions for PLD and RBD segments showing significant overlap. In the strong interaction regime where PLD-PLD interactions dominate, we again observe core-shell structures. However, unlike the spherical disordered, core-shell structures observed when the two interactions are less than 0.5 kT, the PLD core (for *ε*_*PLD*−*PLD*_ *=* 0.6 *kT*) here shows fibril-like nematic ordering. Differential interaction strengths can thereby result in core-shell structures, even for assemblies that are made up of only one type of protein.

**Figure 6:**
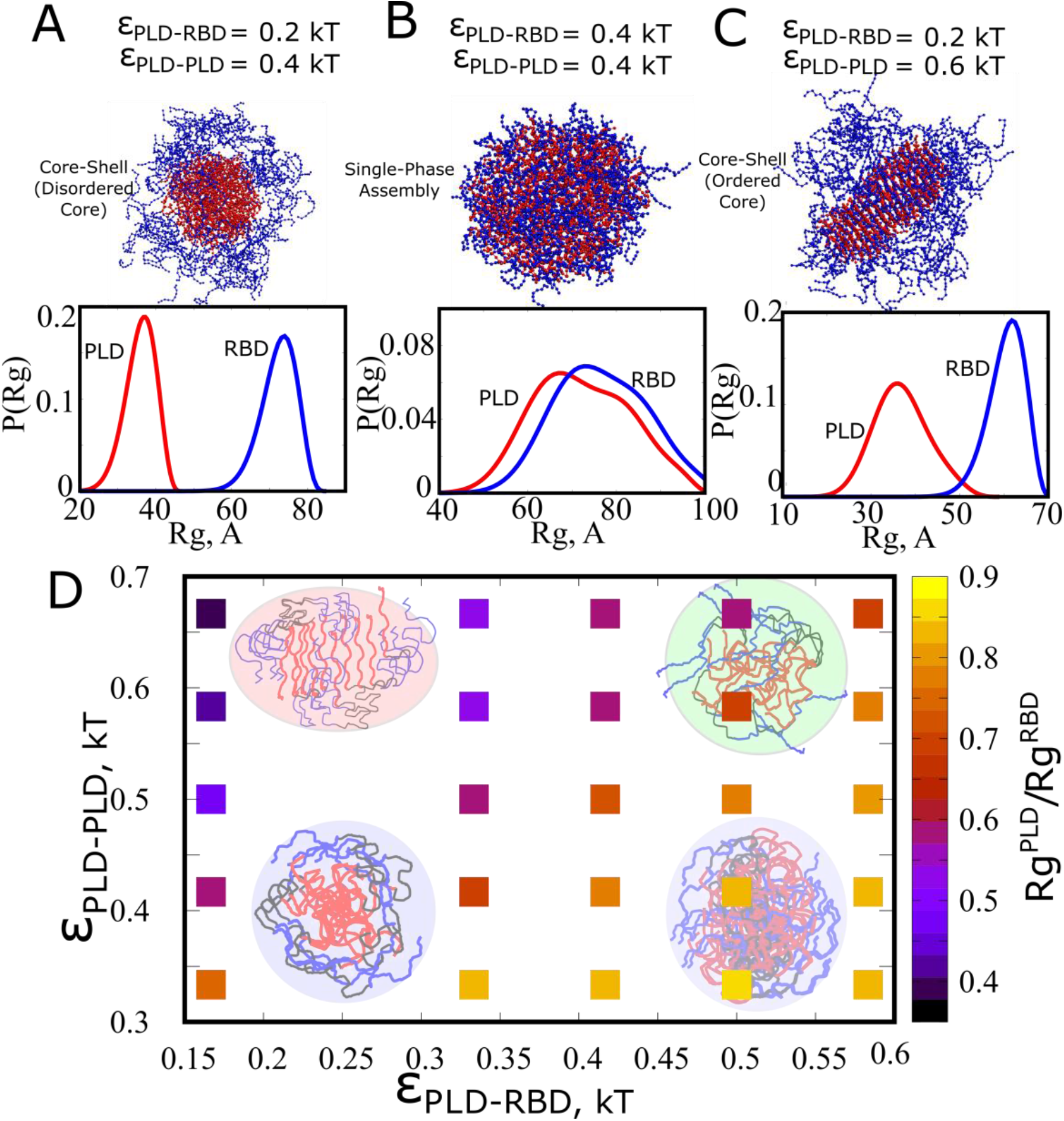
Different architectures of the FUS assemblies. A) For the limit of weak interactions when *ε*_*PLD*−*PLD*_ > *ε*_*PLD*−*RBD*_, we observe disordered structures with the PLDs (red beads) phase separating from the RBDs (blue bead). B) As *ε*_*PLD*−*PLD*_ and *ε*_*PLD*−*RBD*_ become comparable, we observe a disordered assembly where the PLD and RBD domains are fully mixed. C) Above a critical interaction strength, when *ε*_*PLD*−*PLD*_ > *ε*_*PLD*−*RBD*_, we observe core shell structures with distinct PLD and RBD phases within the assembly. Unlike the disordered core-shell structures for weak interactions, the PLD core here shows a high degree of orientational order. The black and purple curves in (A),(B) and (C) show the distribution of radius of gyration of PLD and RBD domains, respectively. Small values of R_g_ indicate a collapsed core while large values indicate that the domains are spread out. When the two distributions do not overlap, it indicates a core-shell structure while overlapping distributions suggest a fully mixed cluster.

### Dramatic slowdown of intra-cluster dynamics upon disorder to order structural transition

Our simulations reveal that the interplay between homo-domain and hetero-domain interactions can result in self-assembled structures with diverse architectures (single vs multiphase, Fig.6) and varying degrees of orientational order. However, aging of droplets and condensates in vivo and in vitro is not only characterized by structural transitions, but also a slow-down in intra-droplet dynamics. In this context, it becomes critical to ascertain whether the sharp disorder-order transitions in structural order and local densities that we observe above a critical ε_PLD-PLD_ are also accompanied by altered intra-cluster dynamics. In order to probe the micro-environment within the FUS clusters, we run long 10-µs long simulations with the self-assembled clusters (localizing ∼100 FUS chains) shown in Fig.3. Using these trajectories, we first plotted the mean square displacements (MSD) of the center of masses of PLD and RBD domains within the cluster (Supplementary Fig.S8 & S9). Here,

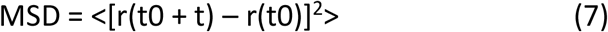

The angular brackets in the equation for MSD indicates an ensemble average across all molecules within a cluster as well as diferent time origins along the trajectory.

We compute the diffusion coefficients (D_PLD_ and D_RBD_) from these MSD curves (by fitting the curves to MSD = 6Dt). In Supplementary Fig. S8, we show the MSD profiles for PLD domains in various scenarios. As evident from Supplementary Fig.S8,S9, the diffusion coefficient for monomeric FUS in solution (∼11 µm^2^/s) shows a 5-fold reduction upon self-assembly into disordered FUS-rich clusters (∼2.5 µm^2^/s). However, while the disordered structures still exhibit diffusive behavior, structures with orientational order (Supplementary Fig. S8, red curve) exhibit flat MSD profiles with two orders of magnitude decrease in D_PLD_. In Fig.7A, we plot D_PLD_ (intracluster diffusion coefficient of PLD domains) as a function of homodomain interaction strengths for a fixed ε_PLD-RBD_ of 0.15 kT. We observed two distinct dynamics regime and a sharp transition between them at ε_PLD-PLD >_ 0.5 kT. As we increase the homo-domain interaction strength from ∼0.5 to ∼0.6 kT, we observe a dramatic – two orders of magnitude - decrease in diffusion coefficients (red curve, Fig.7A, Supplementary Fig.S8), suggesting that the PLD domains exhibit solid-like dynamics in this regime (also evident from the flat MSD profiles in Supplementary Fig.S8 and S8). Interestingly, this sharp transition in diffusion coefficients is concomitant with a sharp increase in structural order (purple curve, Fig.7A) suggesting that solid-like intra-droplet dynamics emerges because of structural ordering. While the disorder-order structural transitions result in an order of magnitude reduction in D_PLD_, the RBD regions show comparatively negligible drop in diffusion coefficients even in highly ordered structures, suggesting that these domains continue to exhibit liquid like dynamics within ordered structures. The spatial heterogeneity in local densities that characterizes PLD-driven assemblies also results in spatially dependent dynamics within the cluster, with a solid ordered core and a liquid-like shell. In Supplementary Fig.S10, we plot detailed phase diagrams that show how the diffusion coefficients of the two domains vary as a function of the homo- and hetero-domain interaction strengths.

**Figure 7:**
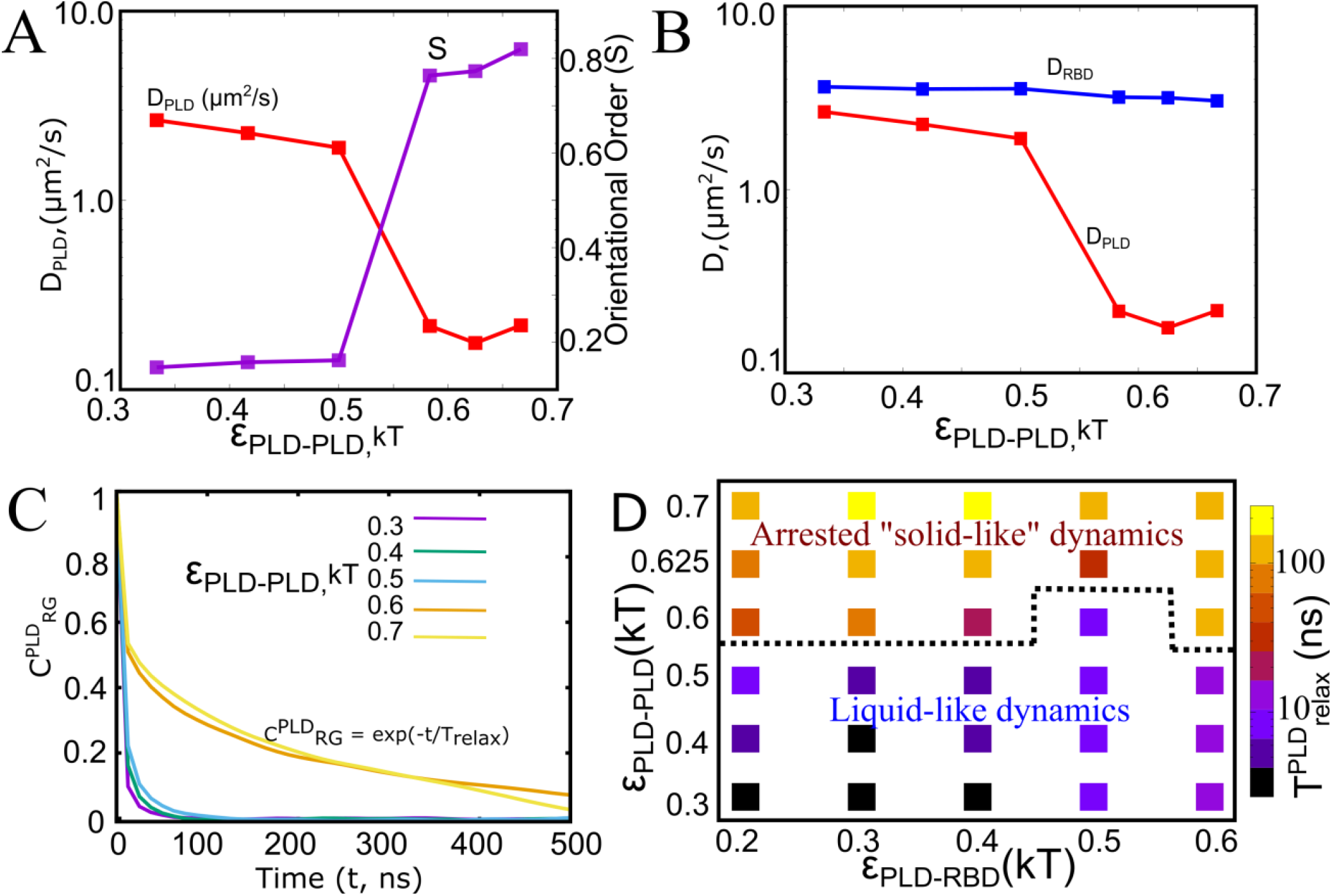
Intra-cluster Dynamics. A) Diffusion coefficients of the PLD domain of polymer chains within the self-assembled cluster, as a function of ε_PLD-PLD_ (red curve, left-hand side Y-axis). As we increase the interaction strength for homodomain interactions above 0.5 kT, we observe a sharp slow-down in intra-droplet dynamics, with an order of magnitude decrease in D_PLD_. This drop in D_PLD_ coincides with a sharp increase in orientational order of PLD domains within the cluster (Purple Curve, right-hand side Y-axis). (B) Diffusion coefficients for PLD (red) and RBD (blue curve) domains within the cluster as a function of ε_PLD-PLD_. At higher values of homodomain interactions, the diffusion coefficient of PLD domains shows an order of magnitude decrease. There is negligible reduction in D_RBD_, in comparison. (C) Normalized Autocorrelation function (C^PLD^_RG_) for the radius of gyration of PLD domains within FUS chains as a function of time. For less viscous intra-cluster environment, the ACF curves decay rapidly to zero, suggesting that the system loses its memory at short timescales. On the other hand, the ACF profiles decay much slower at stronger values of ε_PLD-PLD_ where we encounter ordered structures. C^PLD^_RG_ = exp(-t/T_relax_) D) Measuring polymer dynamics as a function of ε_PLD-PLD_ and ε_PLD-RBD_. T_relax_ is the timescale that defines the rate at which autocorrelation function decays (C^PLD^_RG_ = exp(-t/T_relax_)). The higher this T_relax_, the slower the dynamics of polymers within the protein rich cluster, and more viscous the intra-cluster environment. Low values of T_relax_, on the other hand, indicate that the system loses its memory rapidly and signifies a relatively less viscous intradroplet environment.

We further computed the autocorrelation function, ACF, (computed from time traces of radius of gyration of FUS chains within the clusters) to study the characteristic timescales at which the system loses its memory^47,48^. The normalized autocorrelation function of the radius of gyration of FUS chains, C_RG,_ can be used to describe the dynamics of polymers within the dense polymer-rich clusters.

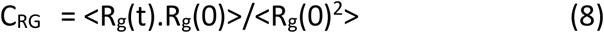

Here, the <> denotes an ensemble average over all the polymer chains that are part of the FUS asssembly as well as using different timesteps as starting point along the trajectory.

As shown in Fig.7C, the ACF profiles decay with time, with faster rate of decay suggesting more liquid-like dynamics while a slower rate of decay is symbolic of more viscous intra-cluster environments. In order to quantify relaxation dynamics, we define an order parameter T_relax_ – T_relax_ is the timescale that defines the rate at which autocorrelation function decays (C^PLD^_RG_ = exp(- t/T_relax_)). T_relax_ allows us to compare the relaxation dynamics of polymer chains within various self-assembled clusters and thereby offers us window into intradroplet dynamics. The higher this T_relax_, the slower the dynamics of polymers within the protein rich cluster, and more viscous the intra-cluster environment. Low values of T_relax_, on the other hand, indicate that the system loses its memory rapidly and signifies a relatively less viscous intradroplet environment. In Figure.7D, we show a phase diagram of polymer relaxation times as a function of the two interaction strengths. Consistent with our MSD profiles, we observed a sharp slowdown in polymer relaxation dynamics at strong ε_PLD-PLD_ suggesting that the regime at which we observe ordered structures is inherently less dynamic. Overall, the different structural phases that we encounter upon tuning the two interaction parameters correspond to distinct dynamic regimes, suggesting that subtle changes in interaction strengths and alterations in interaction networks can result in arrested droplet dynamics.

### Slowdown in intra-cluster polymer dynamics results in altered inter-cluster coalescence behavior

Disorder-order structural transitions result in altered intra-cluster dynamics of polymer chains, with a marked slowdown in polymer relaxation times and diffusion coefficients in the regime corresponding to the ordered phase (Fig.7). Does this altered intra-cluster environment result in altered coalescence of large assemblies? This question is highly relevant in context of biomolecular assemblies where the multi-droplet state has often been observed to be long living (at biologically relevant timescales). Here, we employ metadynamics simulations to probe whether assemblies with varying structural order, formed under different regimes of ε_PLD-PLD_ and ε_PLD-RBD_ have altered coalescence behavior. To address this question, we start off with two large droplets of 100-chains each, and run metadynamics simulations with a biasing potential along a collective variable that captures coalescence -- D_COM._ Here, the collective variable D_COM_ refers to the distance between the center of masses of the two clusters. Previous simulations with system sizes of 50-200 chains have shown that these droplet sizes can recapitulate equilibrium and dynamic quantities in FUS droplets^22^. Large values of D_COM_ (>80 A) correspond to a state where the clusters are not in contact with each other while short distances (< 50 A) suggest that the clusters have undergone complete coalescence. Intermediate values of D_COM_ indicate that the clusters are in contact, without undergoing complete coalescence.

The free energy profiles along this coalescence reaction coordinate – D_COM_ – suggest that disordered, single-phase homogenous condensates (Fig.8A) do not encounter large barriers for coalescence, with a distinct minimum at ∼20 A showing that the two clusters undergo complete mixing to form a single large cluster. Similarly, even core-shell structures formed in the limit of weak interactions show complete coalescence (Supplementary Fig.S11). Strikingly, the partially ordered structures formed in the regime of strong interactions (ε_PLD-PLD_ = 0.7 kT & ε_PLD-RBD_ = 0.6 kT), do not show a free energy minimum corresponding to the coalesced states (Fig.8B). Conversely, due to the strong involvement of both domains in inter-chain contacts, these clusters are uniformly dense (Fig.5C,D & Supplementary Fig.S7). A relatively higher packing density (Fig.5D) of charged domains in this regime (Supplementary Fig.S7) results in an electrostatically-driven emulsification of these droplets. This is consistent with previous studies that suggest that surface charge density can result in a enhanced emulsion stability in biomolecular condensates^26^. The droplets with an ordered PLD core and a less dense, liquid-like RBD shell (Fig.8C) display a free energy minimum for intermediate values of D_COM_. In this regime, the two droplets combine with the ordered ‘fibril-like’ cores joining end to end. However, unlike the partially ordered structures (Fig.8B), in this regime (ε_PLD-PLD_ = 0.6 kT & ε_PLD-RBD_ = 0.2 kT), these structures have a 6-fold lower packing local density (Fig.5D) of charged RBDs, thereby allowing the liquid-like shells of the clusters to penetrate each other. However, the solid-like packing of PLD cores in these structures act as a barrier against complete coalescence. Overall, the altered intra-cluster dynamics of polymers in the various interaction regimes (Fig.7) results in significantly altered coalescence behavior for large clusters. These results suggest that phase transitions that result in a slowdown of intra-cluster dynamics could result in arrested coalescence behavior of biomolecular condensates.

**Figure 8.**
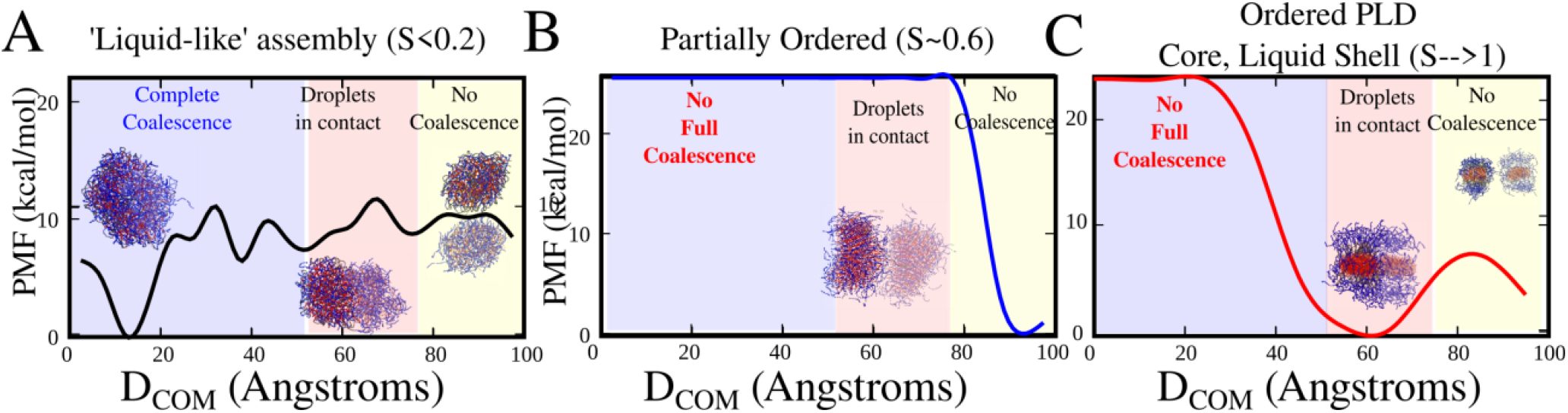
Free-energy profiles for coalescence of large clusters. Free energy profiles for coalescence of two large protein assemblies with different structures, formed at different regimes of ε_PLD-PLD_ and ε_PLD-RBD._ A) Homogenous, disordered droplets (ε_PLD-PLD_ & ε_PLD-RBD_ = 0.2 kT) undergo complete coalescence, with a minima that corresponds to the fully coalesced state. B) Partially ordered structures formed in the limit of strong PLD-PLD and PLD-RBD interactions (εPLD-PLD = 0.7 kT & εPLD-RBD = 0.6 kT) do not coalesce into one large cluster, and remain as individual droplets. C) Assemblies with an ordered PLD-core and a disordered RBD-shell (ε_PLD-PLD_ = 0.2 kT & ε_PLD-RBD_ = 0.6 kT) assemble into one large assembly where the solid cores join end to end without complete coalescence. The order parameter used to track coalescence is the distance between the center of mass of all chains that are members of the first droplet (initially) with those that are part of the second droplet (initially).

### Ordered structures can be seeded by strong non-specific interactions with other scaffold molecules

The contact ratios (Fig.2A), orientational order (Fig.2B and Fig.3) and peak intra-cluster densities (Fig.5) suggest that disordered-ordered transition in FUS clusters is mediated primarily by PLD-driven interactions. Further, the PLD switches from collapsed configuration to an extended ordered configuration above a critical PLD-PLD interaction strength. Therefore, could interactions with domains or regions from other biopolymers also influence ordering within FUS proteins? To address this question we performed simulations with a 200-bead long, homopolymer mimicking RNA molecules that localize within RNP condensates. We first fix the interaction strengths for RNA and RBD beads (*ε*_*RBD*−*RNA*_) to 0.7 kT and then systematically vary the strength of promiscuous RNA binding to the PLD (*ε*_*PLD*−*RNA*_). The strength of *ε*_*PLD*−*PLD*_ and *ε*_*PLD*−*RBD*_ was set to 0.5 kT, a scenario that results in partially ordered structures in the absence of RNA. The ratio of protein to RNA molecules was initially set at 3:1. In Fig.9, we show how RNA-PLD interactions can influence the extent of order in the RNA-protein condensates. In the limit of weak, RNA-PLD interactions, the RNA interactions are extremely specific, binding to the RBD alone, and the system displays partial ordering in PLD domains, akin to the scenario with no RNA molecules. As we increase the strength of non-specific RNA binding to the PLD from 0.1 to 0.3 kT, we see an initial decrease in order because RNA chains compete with PLD domains from other chains for PLD binding (reduction in contact ratio, Fig.9 green curve). However, a further increase in *ε*_*PLD*−*RNA*_ results in an increase in orientational order within these structures, with the PLD and the positively charged RBD showing alignment along the negatively charged RNA backbone.

**Figure 9:**
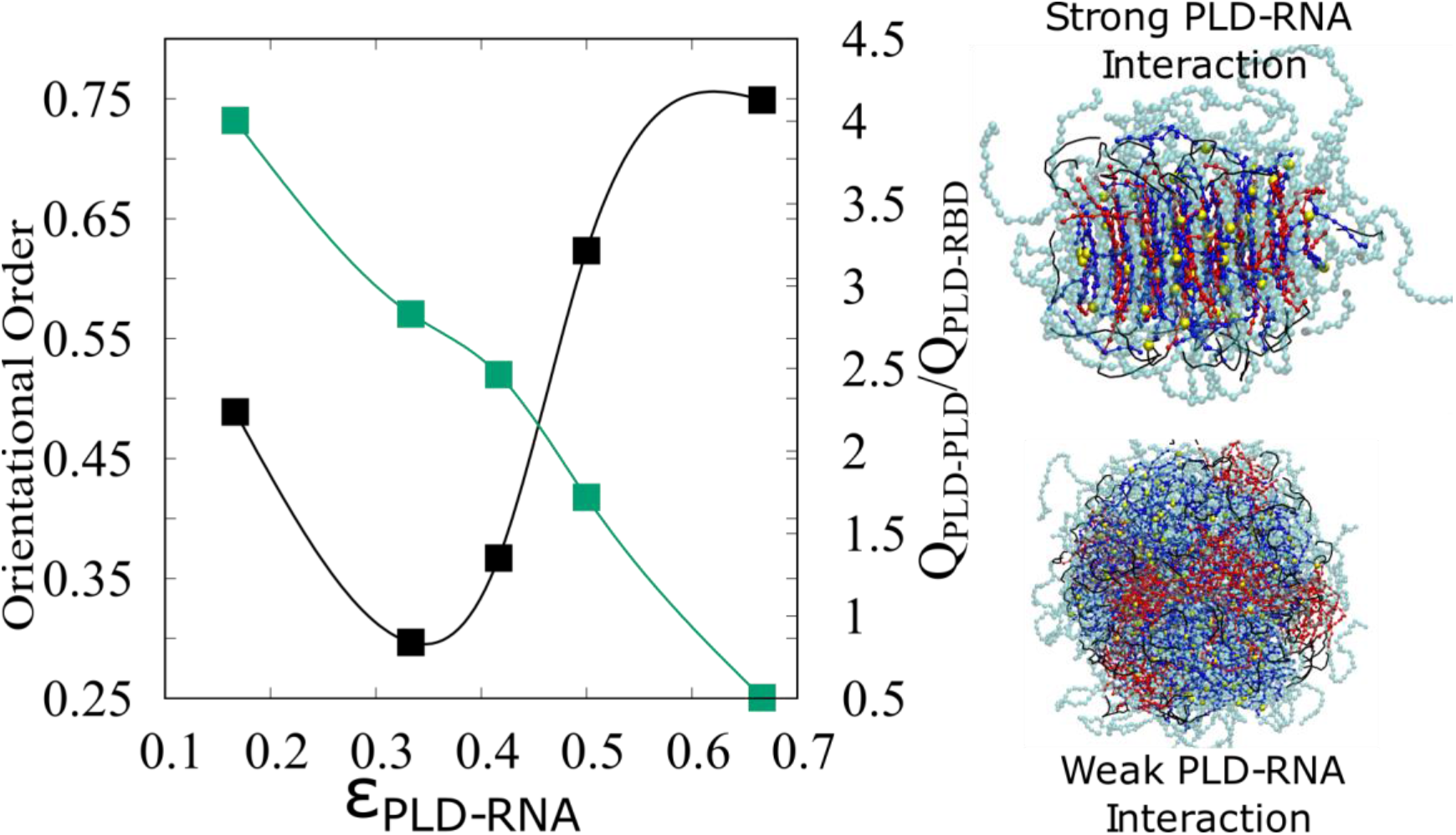
Ordering of FUS-RNA condensates in response to varying strengths of RNA-PLD binding. An initial increase in *ε*_*PLD*−*RNA*_ results in a decrease in orientational order and therefore disordered FUR-RNA assemblies (black curve). A futher increase in *ε*_*PLD*−*RNA*_, however, results in an increase in orientational order, with the PLD domains aligning themselves along the RNA spline. The intermolecular contact ratios (green curve) confirm that the order within the clusters is a result of RNA-PLD contacts and not PLD-PLD contacts.

We further probed the effect of RNA on on structural order in a broad range of RNA concentrations, for different regimes of protein-protein (*ε*_*PLD*−*PLD*_ and *ε*_*PLD*−*RBD*_) and protein-RNA interactions (*ε*_*PLD*−*RNA*_). When protein protein interactions are weak (Fig. 10A, *ε*_*PLD*−*PLD*_ and *ε*_*PLD*−*RBD*_ = 0.4 kT), the structures that form in the absence of RNA (RNA/protein → 0 in Fig.10A and B) are disordered. In this regime, when RNA-protein interactions are weak (*ε*_*PLD*−*RNA*_ *=* 0.2 *kT*, Fig.10A) the RNA-protein assemblies remain disordered for all RNA concentrations under study. However, as RNA binding becomes stronger (*ε*_*PLD*−*RNA*_ *=* 0.4 *kT*, Fig.10B), we observe ordered structures for large RNA concentrations (Fig.10B) in the regime of weak protein-protein interactions. Therefore, for weak protein-protein and strong RNA-protein interactions, RNA molecules can scaffold ordered structures.

**Figure 10:**
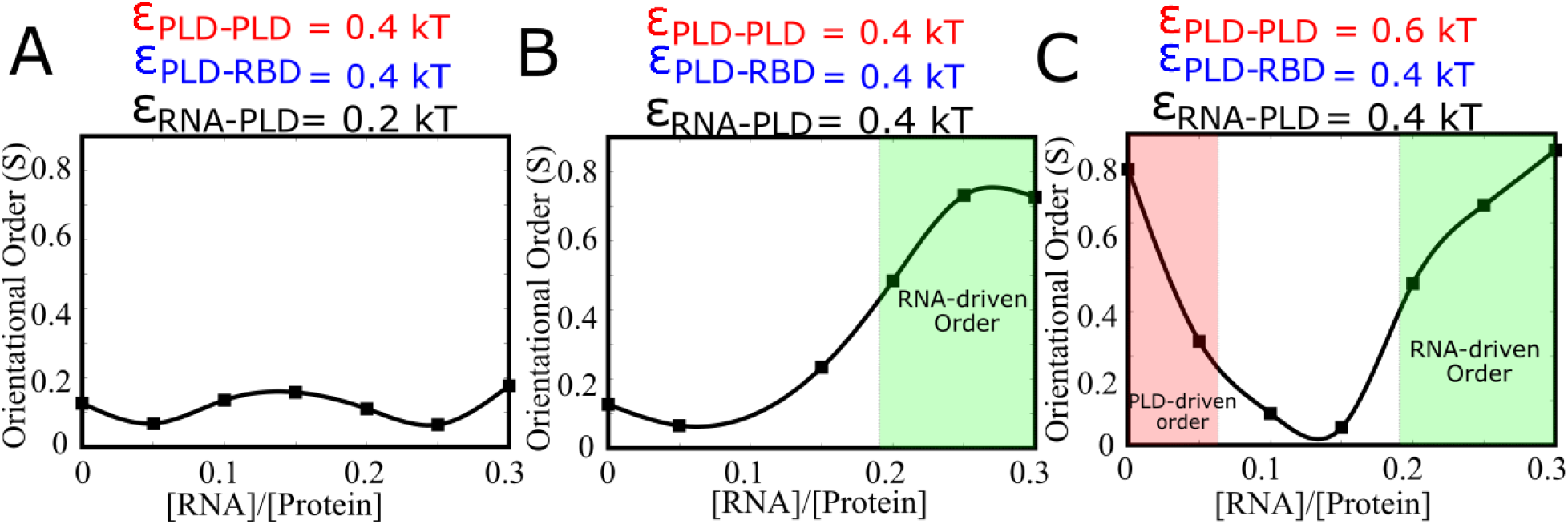
Ordering in FUS-RNA condensates, at varying RNA concentrations. A) When both protein-protein and protein-RNA interactions are weak, *ε*_*PLD*−*PLD*_ and *ε*_*PLD*−*RBD*_ < 0.5 kT and *ε*_*RNA*−*PLD*_ *=* 0.2 *kT*, the ensuing structures remain disordered throught the concentration range. B) As we increase RNA-protein interaction, *ε*_*RNA*−*PLD*_ *=* 0.4 *kT*, we see an RNA-driven ordering of clusters at high [RNA/Protein] ratios. C) When PLD-PLD interactions are strong, the structures that form in the absence of RNA are ordered. As we introduce RNA to the mixture, in this regime of interactions, there is an initial decrease in order. A further increase in RNA concentrations results in ordered RNA-protein clusters, with nematic order as a result of strong RNA-protein interactions.. The solid lines are guide to the eye.

When PLD-PLD interactions are strong (Fig.10C), the structures observed in the absence of RNA display PLD-driven nematic order (Fig.10C, RNA/protein → 0 & Supplementary Fig.S12A). For an initial increase in RNA/protein ratio, we see a decrease in nematic order, due to a competition between RNA and PLD domains for PLD-PLD binding. A further increase in RNA/protein ratio results in an increase in nematic order (Fig.10C). However, the ordering in this regime is a result of RNA-protein interactions, with the RNA forming a backbone around which the ordered structures get stabilized (Fig.10C & Supplementary Fig.S12B).

Therefore, the RNP-clusters exhibit liquid-like, disordered structures only when the strengthj of RNA-protein interactions is within a narrow range. On the other hand, interactions between the PLD domains of different, non-FUS proteins (stronger than a critical *ε*_*PLD*−*PLD*_) could also result in ordered structure formation making this an accessible self-assembled geometry for RBPs with intrinsically disordered regions.

## Discussion

The role of PLDs as key drivers of phase separation can be ascertained by a strong positive correlation between the number of Tyrosine residues in the PLD of FUS family proteins and their measured saturation concentrations for LLPS^16^. While the low-complexity, prion-like domain within FUS can phase-separate on its own, the threshold concentrations for LLPS of PLD alone are higher than that of the full-length protein, suggesting that a hierarchy of interactions could drive self-assembly^17^. PLD domains, on their own can either exist in a disordered droplet like state or an ordered, cross-*β* rich amyloid-fibrillar state. This gives rise to an interesting paradox – while the low complexity domains are crucial drivers of phase separation, a high density of PLDs within the condensed phase makes these structures prone to fibril formation. Indeed, liquid-solid transitions have been widely reported in vitro and upon prolonged stress in vivo^29,46,49^. While the phenomenon of droplet aging and liquid-solid transition is ubiquitous in biological LLPS, its molecular underpinnings are largely unclear. In this study, we study the structural transitions in FUS assemblies within a biologically relevant range of interaction strengths. Are ordered and disordered structural phases characterized by distinct interaction networks? Are morphological transitions accompanied by large changes in intra-droplet densities? Can post-translational modifications act as a barrier to ordered structure formation? To address these questions, we employ a coarse-grained, minimal representation of the multi-domain FUS protein with few interaction parameters in order to understand how the interplay between homo- and hetero-domain interactions could shape the structure of the FUS-rich clusters.

### Disorder-order transition occurs within a biologically relevant regime of interaction parameters and are sensitive to mutations and modifications

Interestingly, the minimal model can effectively capture ordered structures in the narrow range of thermally-relevant interaction strengths (*ε*_*PLD*−*PLD*_ & *ε*_*PLD*−*RBD*_ in the range of 0.1 to 0.7 kT). Our simulations reveal that the disorder-order transition is accompanied by a switch in the predominant interaction network that stabilizes the self-assembled structures. While homo- and hetero-domain interactions are both equally likely in single-phase, disordered assemblies (Fig.3 & Fig.6), the ordered structures are significantly enriched in PLD-PLD interactions (Fig.2A, 3). Crucially, we observe that this transition from the disordered to ordered self-assembled state has the hallmarks of a highly cooperative first order phase transition (Fig.2B). A small change in homodomain interaction strengths (≈ 0.1 kT per bead, Fig.2B,C and Fig.3) could result in dramatically altered structures. Mutations of single or small numbers of PLD residues, resulting in altered strength of interactions and/or interaction network could therefore cause disorder-order transitions in FUS protein droplets. Further evidence that ordered structures are dependent on PLD-PLD interactins comes from simulations with phosphorylated PLDs that localize a high density of negative charges. Consistent with experimental findings^29^, phosphorylation resulted in down-regulation of PLD-PLD contacts resulting in structures with low nematic order. Phosphorylation not only resulted in disordered structures (Fig.4) but also lower local densities (Fig.5B) of FUS PLD compared to assemblies composed of unmodified FUS. Also, ordered structures exhibited several-fold higher peak local densities compared to disordered phases, suggesting that these structures could also have vastly different material properties

### The origins of structural order in the semi-flexible polymer model

The disorder-order transition in our simulations is reminiscent of the isotropic-nematic transitions in liquid-crystals that occur at low temperatures or high density. However, unlike liquid-crystals, the disorder-order transition in our model occurs in the semi-flexible limit (*L*_*PLD*_ and *L*_*RBD*_ >> *L*_*p*_) as opposed to the rod-like limit, indicating that the ordered phases are formed to maximize favorable PLD-PLD interactions at the expense of competing, less-favorable PLD-RBD and RBD-RBD interaction. Interaction-driven ordering of polymer self-assemblies has been previously reported for homopolymer chains^35^. This mechanism is also observed in polymer crystallization where higher density lamellae consisting of ordered chains intertwine with disordered regions^50,51^. Di-block copolymers have previously been known to assemble into structures with liquid-crystalline order within a condensed phase^52^. In this context, the multi-domain architecture of FUS, with the uncharged PLD and the positively charged RBD, is analogous to a di-block polymer. Inter-chain interactions between biopolymers like proteins can therefore play the role of density modulators within the condensed state. When the strength of these interactions exceeds a critical limit, it results in sharp density transitions within the self-assembled phase (Fig.5C) reminiscent of an Onsager-like isotropic-nematic transition^53^. Unlike the Onsager-like model where the liquid crystals are rod-like molecules that exclude volume, the polymer chains in our simulations are self-associative and semi-flexible. Above a critical strength of interactions, the loss of entropy (loss of flexibility) from ordering is compensated for by the gain of enthalpy by strong inter-chain interactions in the densely packed ordered state where resulting sheet like ordering provides higher local density and less defects. While structural order could be a result of either homotypic (PLD-PLD) or heterotypic interactions (with RNA or with RBD) stronger than a critical limit, the high density of positive charges on the RBD of FUS does not allow local densities large enough to promote ordering (Fig.5 & Supplementary Fig.S7). Demixing of PLDs and RBDs within the condensed phase in order to maximize PLD-PLD interactions and significantly increase local packing density is therefore the most plausible mechanism for ordering within diblock co-polymer condensates such as FUS. Given that subtle changes to interaction strengths result in dramatically altered order, such transitions could be a response to even slight changes in interaction parameters such as point mutations or solvent conditions..

### Disorder-order transitions are liquid-solid transitions

Strikingly, the sharp density (and structural) transitions resulting from tuning the interaction parameters are also accompanied by dramatically altered intra-droplet dynamics, as shown by the intra droplet diffusion coefficients and polymer relaxation times (Fig.7 and Supplementary Fig.S8-S10). These results reveal that different structural phases also exhibit distinct material properties ranging from liquid-like (fast relaxation dynamics) to irreversible, trapped, solid-like (slow relaxation dynamics). While liquid-like disordered structures have diffusion coefficient 5-times lower than the dilute phase, the ordered structures show a 50-fold reduction in diffusion coefficients (Supplementary Fig.S8-10). This vastly altered dynamics is highly relevant to biological condensates where an irreversible phase transition to the solid-like state is a signature of abberant behavior^54^. Indeed, FUS condensates have been identified to span several different material states, with ALS mutations accelerating their transition to the irreversible, solid-like state that results in impaired RNP-granule functions^24,54^. Our results therefore provide a minimal, mechanistic framework that link structural properties of the protein assembly to its material state.

## Testable Predictions

A key feature of the FUS protein (and other RNA-binding proteins of the FUS family) is that the two domains – the PLD and the RBD – have distinct amino-acid compositions. The PLD is rich in Tyrosine residues while the disordered regions of the RBD localize Gly and Arg residues. Previous studies have shown that the number of Tyrosines in the PLD and Arginines in the RBD contribute to the multivalency of FUS^17^. Our findings emphasize that the structural and material fate of the FUS assembly is decided by the interaction network that stabilizes the condensed phase. The interaction parameters in our simulations, *ε*_*PLD*−*PLD*_ and *ε*_*PLD*−*RBD*_, can be tuned by sequence modification (mutations, phosphorylation of PLD, methylation of RBD) or by varying solvent conditions (salt concentration, pH, addition of RNA). Here, we propose some experiments that could be used to test the key findings from our simulations.

a. Strengthening PLD-PLD interactions and weakening PLD-RBD interactions results in an increased propensity to self-assemble into ordered aggregates stabilized by PLD-PLD interactions. In this context, experiments that selectively weaken pi-cation interactions between Tyr and Arg residues of the PLD and RBD domains could result in a PLD-dominant condensed phase. Therefore, a systematic experimental study of FUS assembly at increasing salt concentrations would be extremely relevant for testing the predictions from our model. Our results suggest that at low salt concentrations, the hetero-domain pi-cation interactions between Tyr and Arg residues would dominate, whereas at high salt, one would expect condensates that are stabilized by homodomain pi-pi interactions between Tyr residues of the PLD. It must, however, be noted that the threshold concentrations for LLPS could be different at different salt concentrations.
b. Similarly, one could also setup experiments wherein hydrophobic interactions between the Tyr residues of PLDs can get selectively disrupted in order to test our predictions of PLD-driven ordering. 1,6-hexanediol, an aliphatic alcohol has previously been employed to probe material properties of phase separated droplets in response to selective disruption of hydrophobic interactions^55^. Our simulations predict that assemblies that form at increased concentrations of such a hydrophobic disruptor (that weakens PLD-PLD interactions) would show a lower likelihood of transitioning to an ordered nematic-like assembly.
c. Our simulations also suggest that subtle changes to the interaction strengths (of the order of 1 kT) could result in dramatically altered ordering within the assembly, confirmed by abrogation of order within the phosphorylated variants. In this context, studying sequence variants wherein the Arg residues of the RBD are substituted to reduce their net interaction propensity with the Tyr residues of the PLD, could also result in a switch in the interaction network. Similarly, we also predict that one could also reduce the propensity to form ordered structures by studying variants with Tyr-substitutions in the PLD.
d. One of the most striking predictions from our model is that the ordered states that are characterized by PLD-PLD interactions are also dynamically solid-like. In this context, studying the material properties of condensates (using FRAP studies) formed under different experimental conditions described above (varying salt, 1,6-hexanediol concentrations) could be really relevant to establish the relationship between structure (determined by the key stabilizing interaction network) and material properties of the droplets.

### Conclusions

The fact that a minimal model lacking residue-level resolution can capture diverse structural phases associated with protein condensates -- single-phase, core-shell disordered and nematically ordered structures -- suggests that the assembly of these structures could follow some generic physical mechanisms. While the current study employed a coarse-grained version of the FUS protein, the general principles for forming ordered, nematic-like structural phases could be applicable to other, related multi-domain RBPs that harbor intrinsically disordered domains (Figure. 11). The observation that a rich diversity of structural phases could exist for a narrow range of interaction strengths suggests that these transitions are subject to tight regulatory control. Any deviation from this tight regulation could result in aberrant phase transitions that are an outcome of subtle changes to the interaction networks.

**Figure 11:**
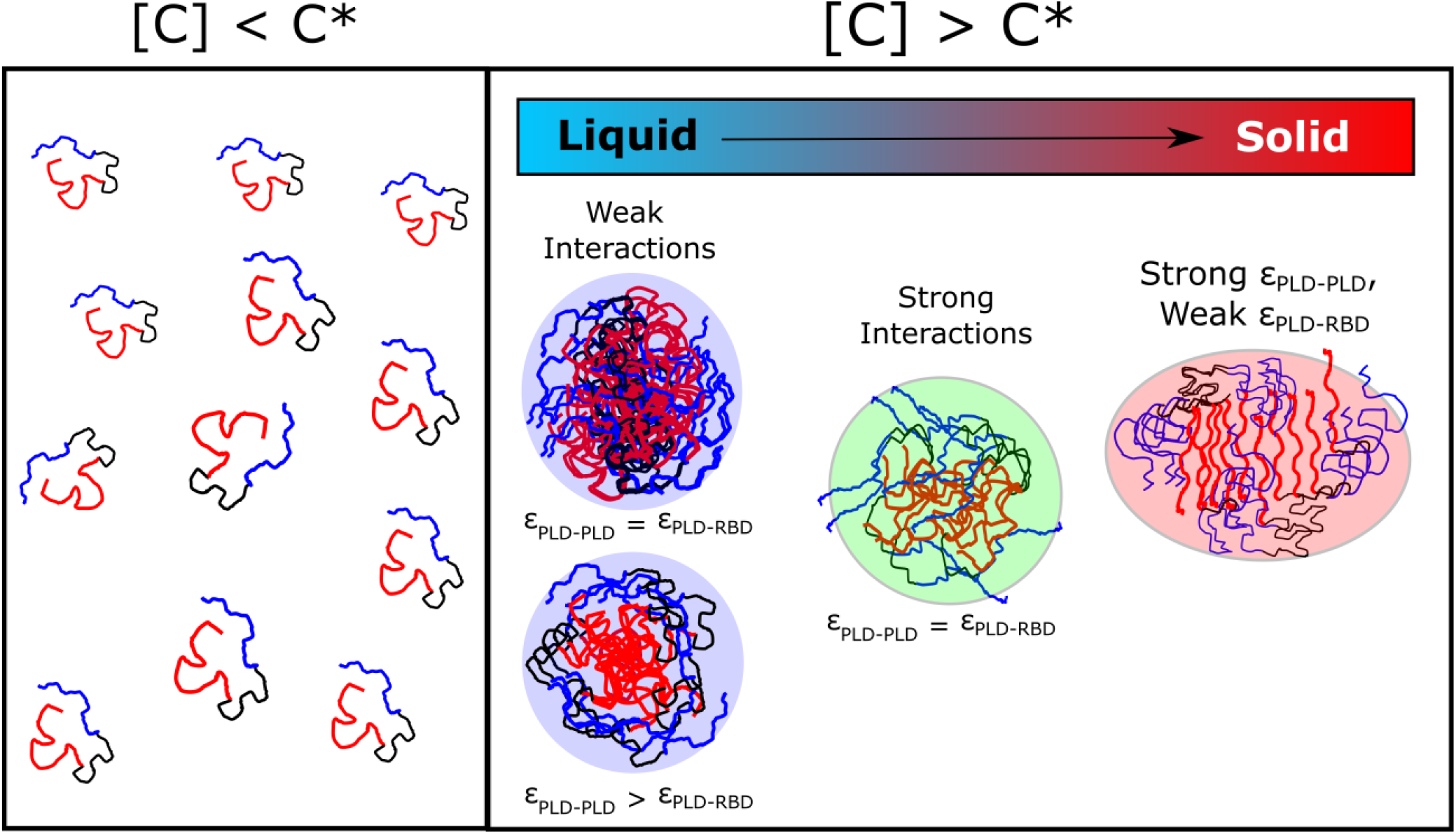
Graphical Summary of the findings. Below a critical concentration C*, the multi-domain RBPs remain in their soluble, fully-mixed state. However, above the critical concentration, these structures could exist in different structural and material phases that are an outcome of the interplay between homodomain (PLD-PLD) and heterodomain (PLD-RBD) interactions.

## Author Contributions

SR was involved in conceptualization of the work, analysis and writing the paper. ES conceptualized the work, analyzed the results and wrote the paper. SR performed the simulations.

