## Supplementary Figures and Tables for "The physics of liquid-to-solid transitions in multi-domain protein condensates"

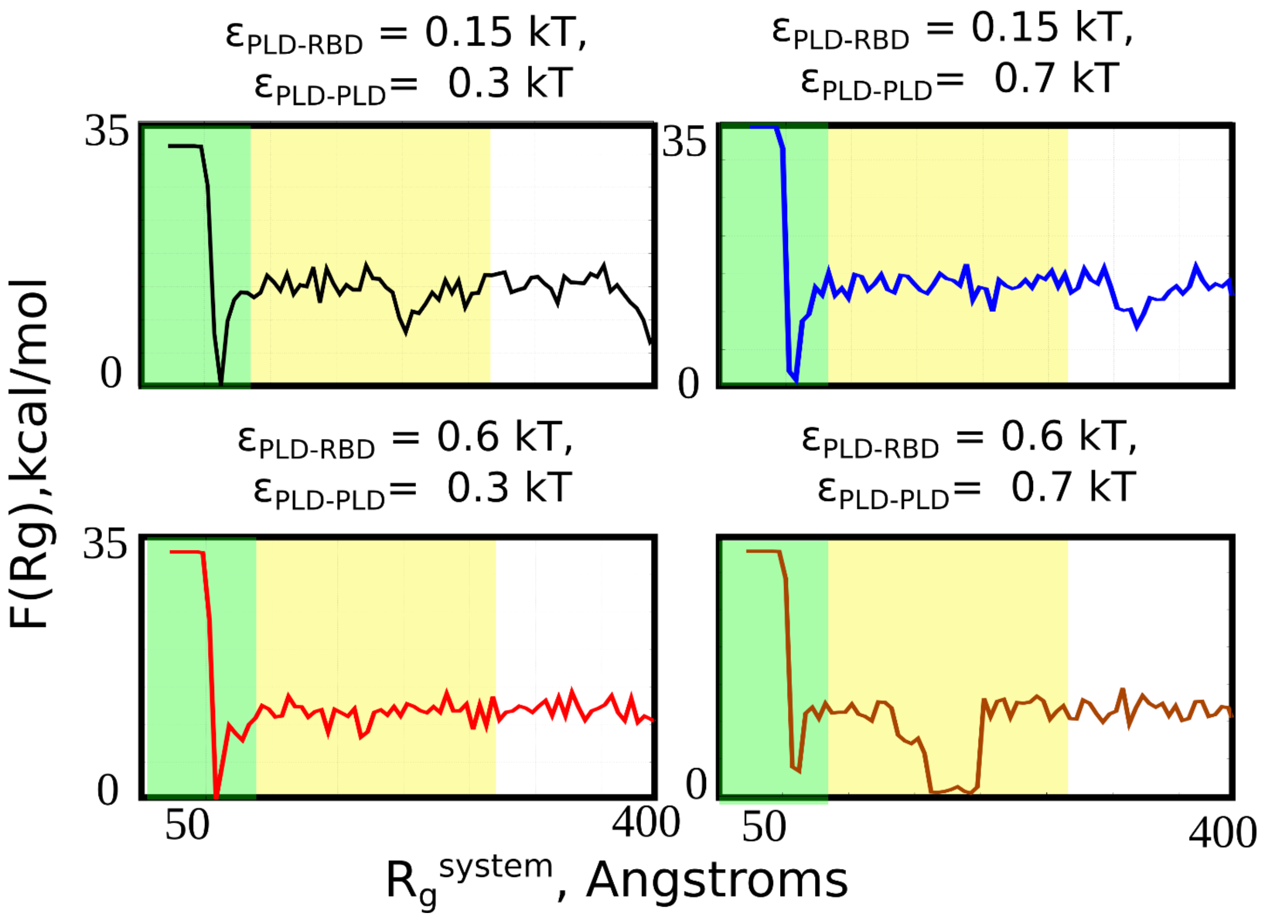


**Figure S1.** Free energy profiles for different combinations of PLD-PLD and PLD-RBD interaction strengths. A sharp minima at ≈ 50 Å (shaded green) is indicative of an equilibrium state corresponding to a single large self-assembled cluster. As $R_g^{system} \rightarrow 400$, the system approaches the fully mixed state.

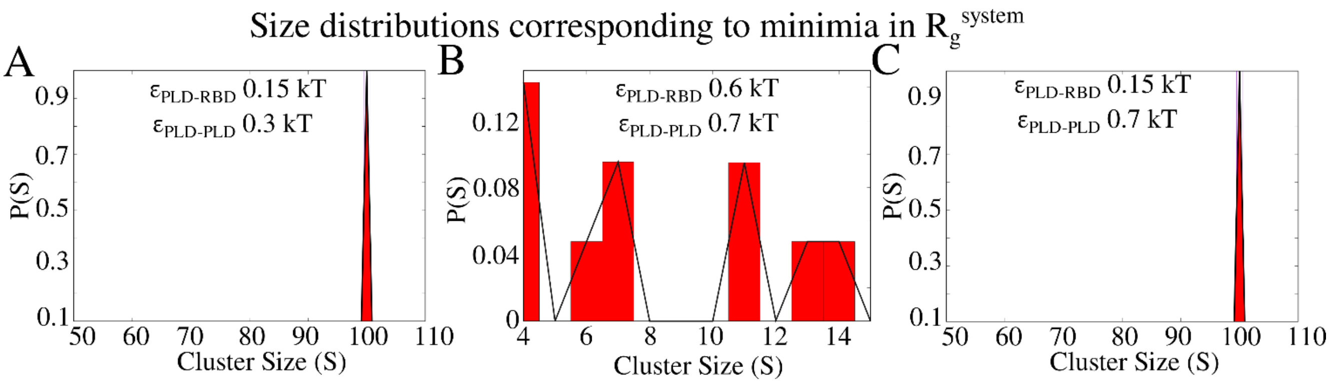


**Figure S2.** Cluster size distributions corresponding to the free energy minimum in metadynamics simulations.

**Free Energy Profiles in Fig.S1 reveal that the condensed phase is favored under different interaction regimes.**

In order to drive the self-assembly of the proteins into a single, large, condensed phase, we employ metadynamics simulations. We use the radius of gyration of the system ($R_g^{system}$ of $N$ protein chains as a collective variable to define the self-assembled state of the system. The free energy profiles were computed along this reaction coordinate, with minima around small values of $R_g^{system}$ indicating a fully assembled equilibrium state. In Supplementary Fig.S1, we demonstrate the free energy profiles for four such scenarios, with varying strengths of hetero-domain ($\epsilon_{PLD-RBD}$) and homodomain ($\epsilon_{PLD-PLD}$) interactions. As evident from Fig.S1, for all the four scenarios, we observe a distinct minima for $R_g^{system}$ ≈ 50 A (shaded green region), a value that corresponds to a state where there is a single large condensed phase. The yellow region in the free energy profiles corresponds to the multi-cluster state, with local minima in this region potentially indicative of kinetic barriers to cluster growth. The free energy profiles suggest that the protein concentrations and the range of interaction strengths chosen for study favor self-assembly of FUS into a large condensate (≈ 100 FUS chains). The metadynamics simulations also reveal that FUS clusters could form in the limit of weak (top panels in Fig.S1) or strong hetero-domain interactions (bottom panels in Fig.S2). Crucially, even when both homotypic interactions ($\epsilon_{PLD-PLD}$) and the heteropypic ones ($\epsilon_{PLD-RBD}$) are relatively weak (0.3 and 0.15 kT respectively), the system tends to assemble into protein-rich clusters.

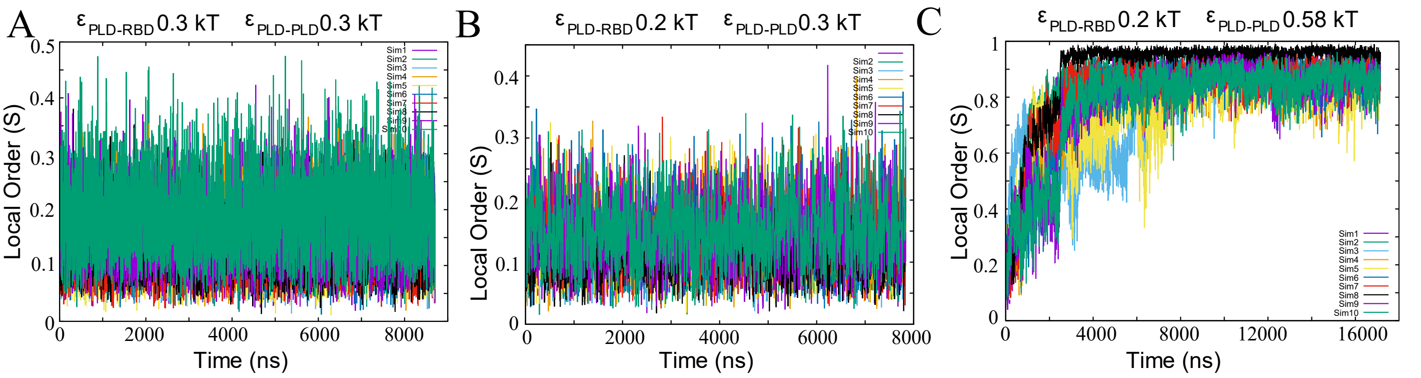


**Figure S3.** Evolution of orientational order parameter (S) for 10-independent realizations of the simulation. The three panels A,B and C, correspond to different regimes of interactions.

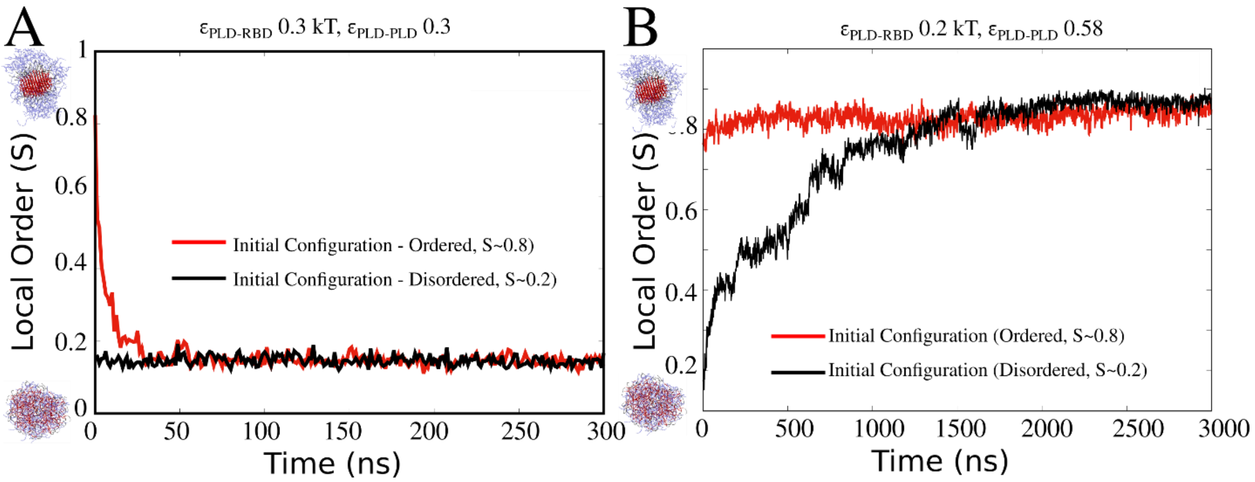


**Figure S4. Convergence of order parameter for simulations with different initial configurations.** Ordered structures (S~0.8) relax to structures with weak order (S~0.2) upon weakening interaction parameters. Conversely, upon increasing strength of PLD-PLD interactions, disordered structures reorganize to form fibril-like order structures at simulation timescale.

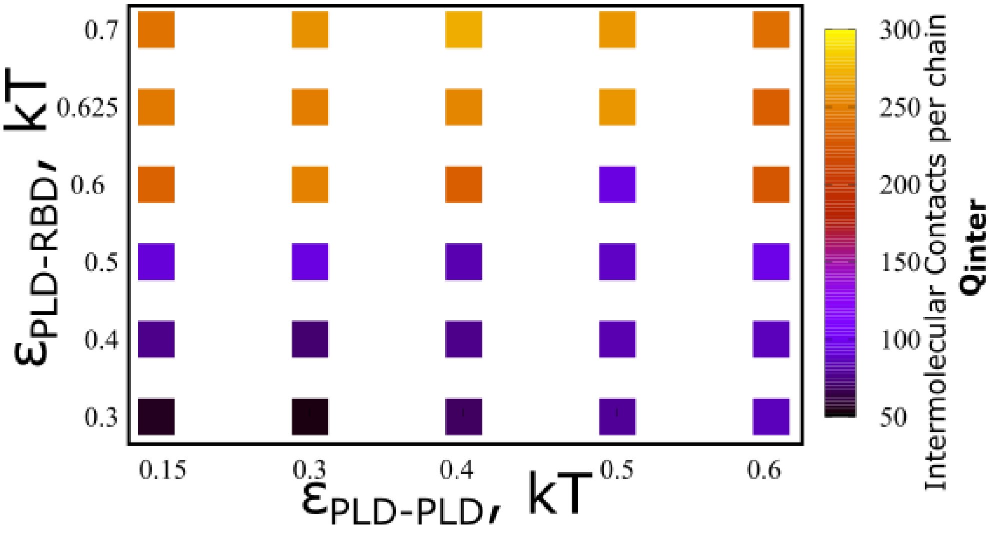


**Figure S5.** Inter-chain contacts per chain, as a function of varying PLD-PLD and PLD-RBD interaction strengths.

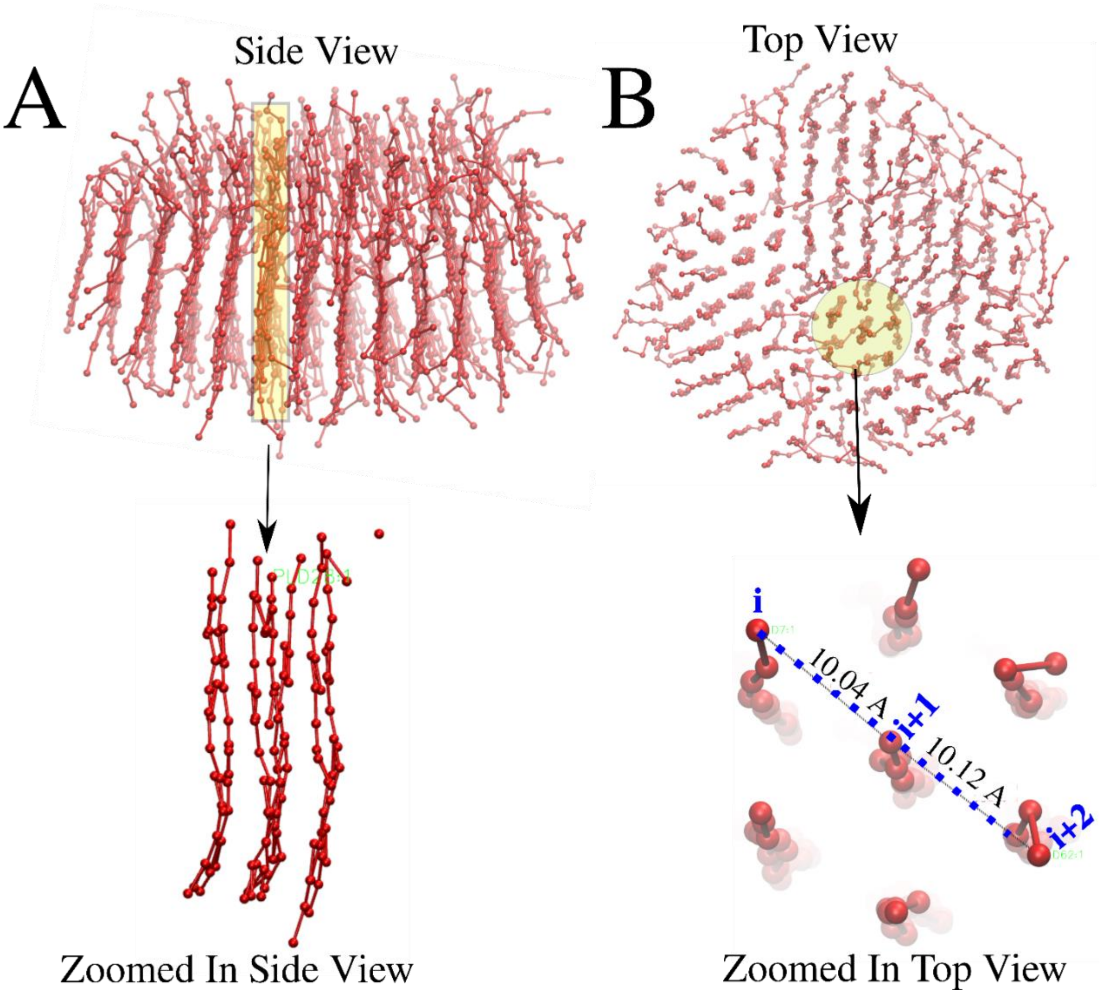


**Figure S6.** Visualization of the ordered PLD core within ordered core-shell structures. Panel (A) shows a side view representing the dense packing of the PLDs with high orientational order where in sheets are stacked alongside. (B) Top view of the orientationally ordered PLD core showing periodic arrangement of PLD chains. The bottom panels in (A) and (B) show a zoomed in version of a local region within the dense core. The lower panel in (B) shows the characteristic spacing between the i,i+1 and i+2th chains within the PLD core.

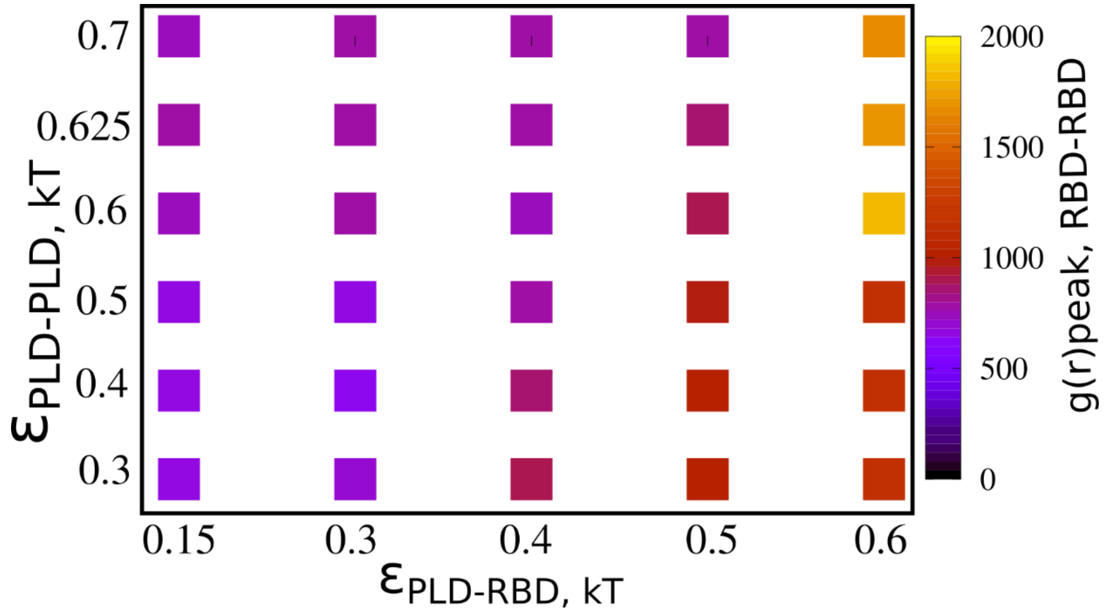


**Figure S7.**Peak values of pair distribution function **g(r),** for RBD-RBD contacts. As the strength of PLD-RBD interactions increases, we see an indirect effect on the peak local density of RBD-RBD interactions.

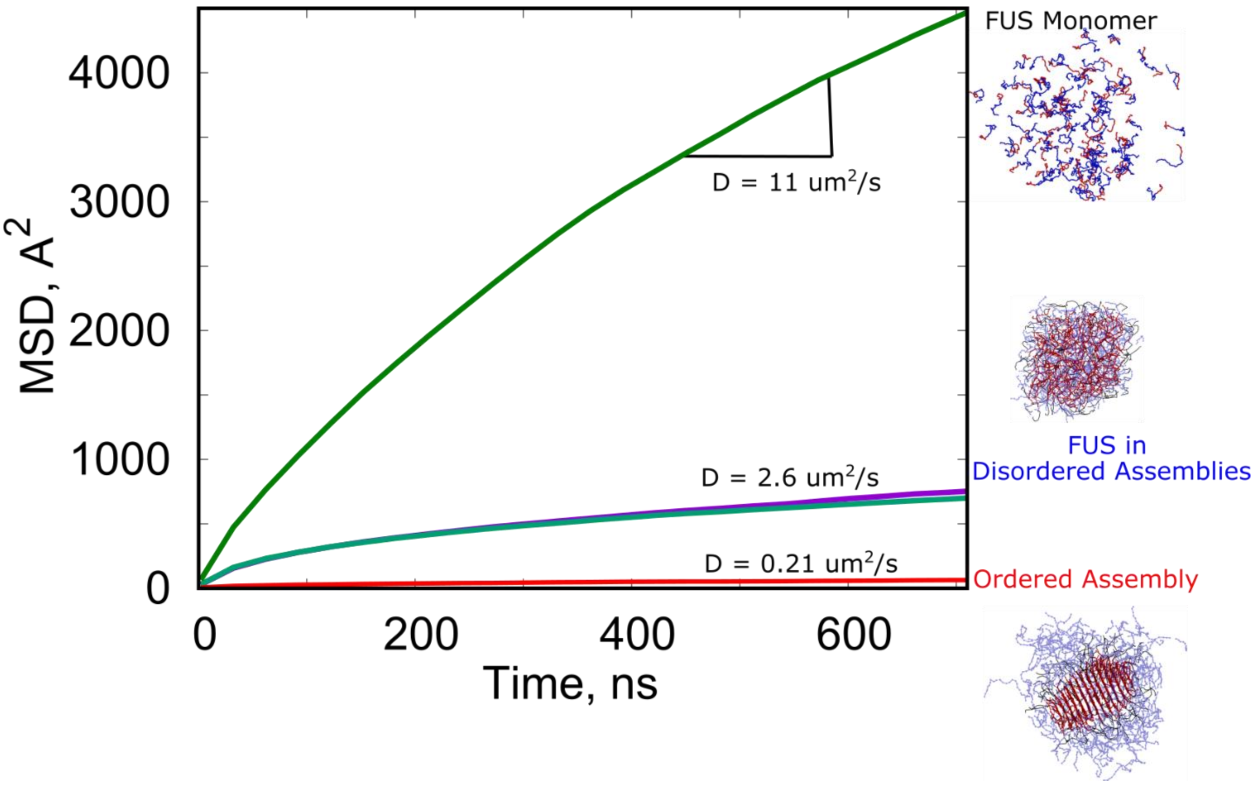

**Figure S8.** Mean square displacement profiles for FUS chains in different scenarios – in the monomeric state (green curve), within single and multi-phase disordered droplets (blue curve) and ordered assemblies (red curve).

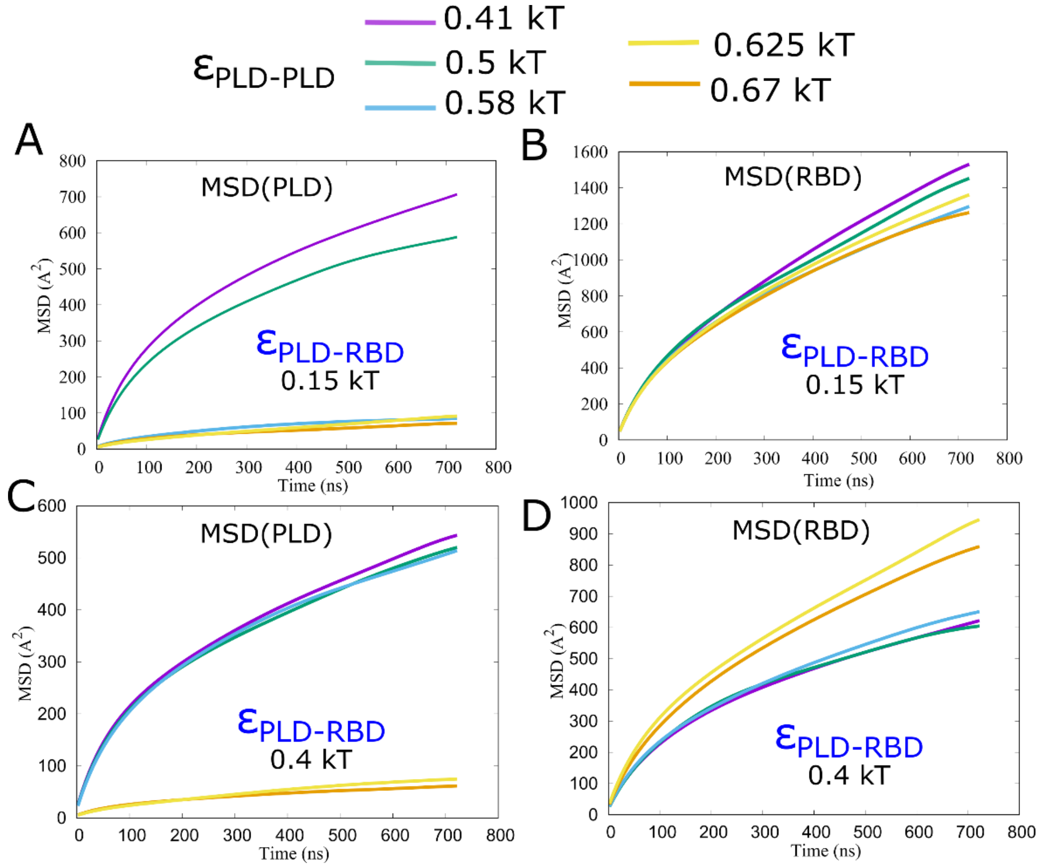


**Figure S9. Mean Square Displacement** profiles for the PLD (A and C) and RBD (B and D) domains, for different strengths of PLD-RBD (0.15 kT in panel A and B) and PLD-PLD interactions (different colored curves in panels A-D).

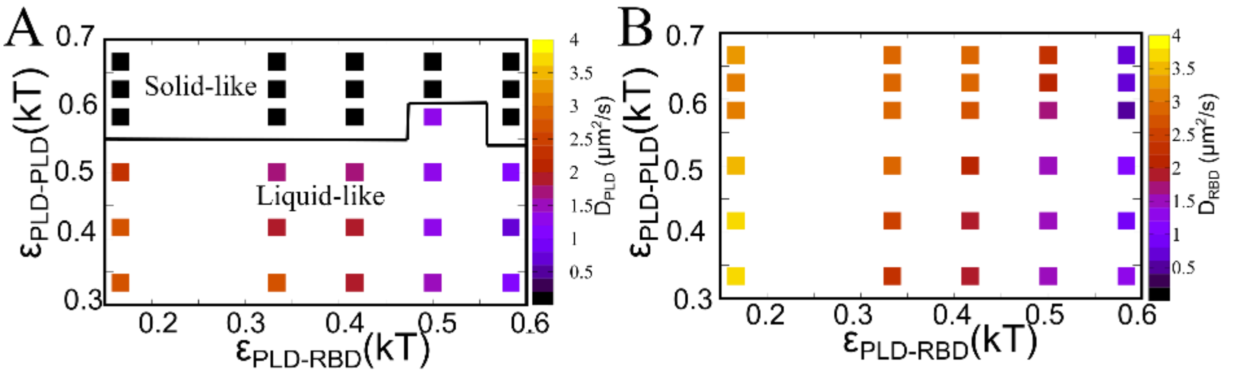


**Figure S10. Diffusion Coefficient phase diagrams.** A) Diffusion coefficient for PLD domains showing dramatically lower values at higher values of PLD-PLD interactions. B) The RBD regions remain liquid like in the whole range of parameters studied.

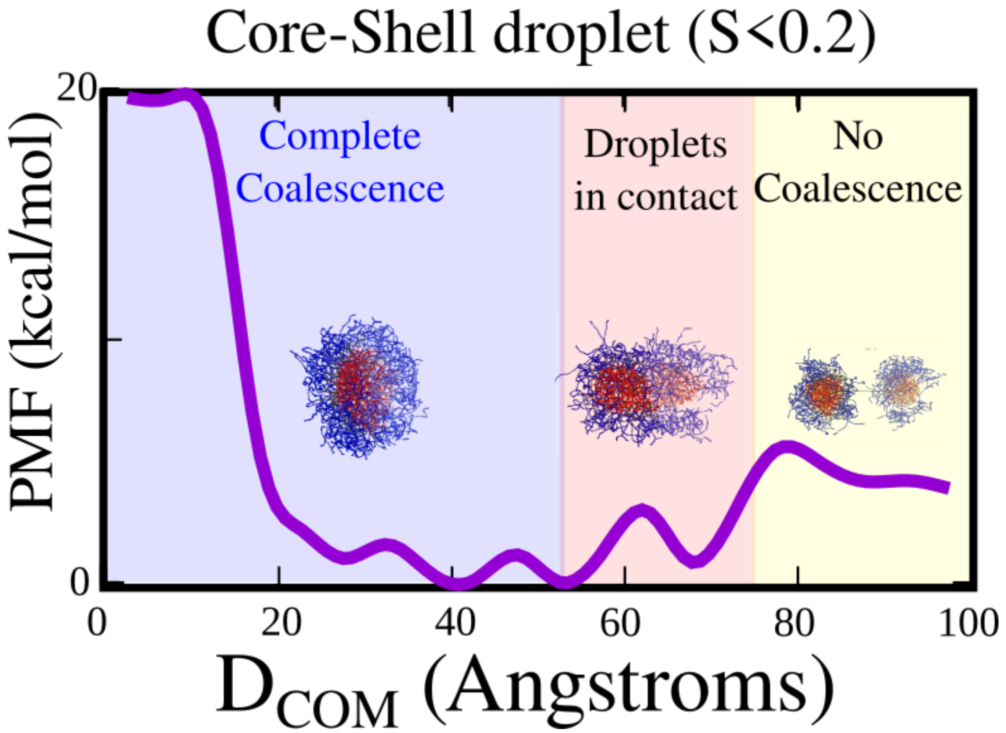


**Fig. S11. Coalescence simulations for Core-shell disordered droplets.** Core-shell, disordered droplets ($\varepsilon_{PLD-PLD}$ = 0.3 kT & $\varepsilon_{PLD-RBD}$ = 0.15 kT) display a minima that corresponds to the fully coalesced state.

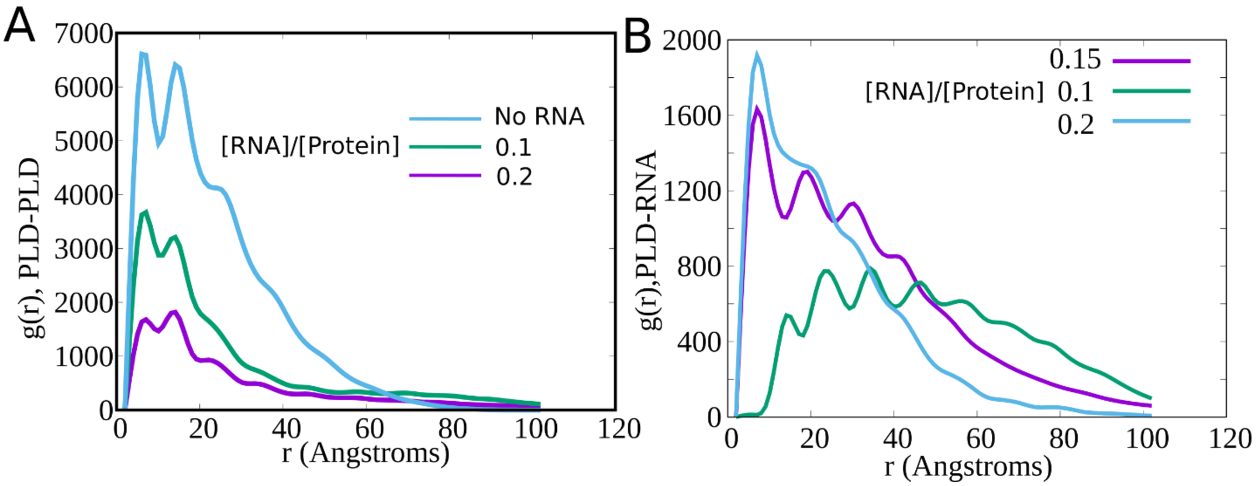


**Figure S12.** Evolution of pair distribution function **g(r),** for increasing concentrations of RNA. (A) g(r) curves for PLD-PLD contacts, and (B) for PLD-RNA contacts.

### Supplementary Information Table S1.

### Human FUS Sequence

```
>sp|P35637|FUS_HUMAN RNA-binding protein FUS OS=Homo sapiens
OX=9606 GN=FUS PE=1 SV=1
MASNDYTQQATQSYGAYPTQPGQGYSQQSSQPYGQQSYSGYSQSTDTSGYGQSSYSSYGQSQN
TGYGTQSTPQGYGSTGGYGSSQSSQSSYGQQSSYPGYGQQPAPSSTSGSYGSSSQSSSYGQPQ
SGSYSQQPSYGGQQQSYGQQQSYNPPQGYGQQNQYNSSSGGGGGGGGGGNYGQDQSSMSSGGG
SGGGYGNQDQSGGGGSGGYGQQDRGGRGRGGSGGGGGGGGGGYNRSSGGYEPRGRGGGRGGRG
GMGGSDRGGFNKFGGPRDQGSRHDSEQDNSDNNTIFVQGLGENVTIESVADYFKQIGIIKTNK
KTGQPMINLYTDRETGKLKGEATVSFDDPPSAKAAIDWFDGKEFSGNPIKVSFATRRADFNRG
GGNGRGGRGRGGPMGRGGYGGGGSGGGGRGGFPSGGGGGGGQQRAGDWKCPNPTCENMNFSWR
NECNQCKAPKPDGPGGGPGGSHMGGNYGDDRRGGRGGYDRGGYRGRGGDRGGFRGGRGGGDRG
GFGPGKMDSRGEHRQDRRERPY
```

### Supplementary Information Table S2.

### Human FUS Phosphosites

| | |
|---|---|
| | S3,T7,T11,T19,S26,S30,S37,S42,S54,S57,S61,T68,T71,S77,T78,S84,S86,S87,S95,S96,S108,S109,S110,S112,S115,S117,S127,S129,S131,S135,S142,S148 |

### Supplementary Information Table S3.

### Pairwise Interaction Parameters

| Notation | Physical Interpretation | Values used in simulation |
|---|---|---|
| $\varepsilon_{PLD-RBD}$ | Strength of LJ-attractive interaction between PLD and RBD beads. *Here, the notation RBD refers to the regions of the RNA-binding domain other than the folded RNA-recognition motif (RRM) and the Zinc-finger motif (ZnF). | Varied in the range of 0.2 to 0.6 kT (pairwise) |
| $\varepsilon_{PLD-PLD}$ | Strength of LJ-attractive interaction between PLD and RBD beads. | Varied in the range of 0.2 to 0.8 kT (pairwise) |
| $\varepsilon_{LIN-LIN}$<br>$\varepsilon_{LIN-RBD}$<br>$\varepsilon_{LIN-PLD}$<br>$\varepsilon_{LIN-RRM}$<br>$\varepsilon_{LIN-ZnF}$<br>$\varepsilon_{RBD-ZnF}$<br>$\varepsilon_{PLD-ZnF}$<br>$\varepsilon_{PLD-RRM}$<br>$\varepsilon_{RRM-ZnF}$<br>$\varepsilon_{ZnF-ZnF}$<br>$\varepsilon_{RRM-RRM}$ | Strength of LJ-attractive interactions involving the flexible linker (LIN) and the folded RNA-recognition motif (RRM) and the Zinc-finger domain (ZnF) | Weakly attractive at 0.1 kT (pairwise). Primarily excluded volume interactions. |

| | | |
|---|---|---|
| $\varepsilon_{RRM\text{-}RNA}$ & $\varepsilon_{ZnF\text{-}RNA}$ | Strength of LJ-attractive interactions between coarse-grained RNA chains and the folded regions – RRM and ZnF – within the RBD. | Attractive interations set to 0.6 kT (pairwise). |
| $\varepsilon_{RBD\text{-}RNA}$ | Strength of LJ-attractive interactions between coarse-grained RNA chains and the disordered regions – labeled RBD – within the RBD. | Attractive interations set to 0.4 kT (pairwise). Additionally, the +vely charged RGG-beads (labeled RBD in the notation) also interact with the negatively charged RNA beads via screened electrostatic interactions. |
| $\varepsilon_{PLD\text{-}RNA}$ | Strength of LJ-attractive interactions between the PLD and RNA. | Varied in the range of 0.1 to 0.7 kT |
| $\varepsilon_{RNA\text{-}RNA}$ | Strength of LJ-attractive interaction between RNA beads | Excluded volume interactions only. RNA beads also harbor a negative charge of -1 per bead. |
| $\sigma_{PLD}$, $\sigma_{RBD}$ and $\sigma_{LIN}$ | Size (radius) of PLD and RBD beads.<br><br>Note: RBD refers to the disordered regions within the RNA binding domain. | Each bead coarse-grains about 6-8 amino acid residues. Diameter of each amino-acid, $\sigma_{aa}$, is about 4.5 A. Therefore, assuming that the 6-8 amino-acid stretches behave like random coils, $N_{aa}^{\Upsilon}\sigma_{aa.}$ Using $\Upsilon$=2/5 for a |

| | | |
|---|---|---|
| | | random coil, we get a bead radius of about 6 Angstrom. Note that ϒ=2/5 was chosen based on the Flory theory for homopolymers in good solvent. |
| $\sigma_{RRM}$, $\sigma_{ZnF}$ | Size (radius) of ZnF and RRM domains | Chosing a scaling exponent of 0.33 for folded domains to model compactness, we get bead sizes of about 8 A (radius) for these regions. |
